# Intron Retention Controls Localization of lncRNAs *PURPL* and *MALAT1* to Promote Cell Proliferation and Migration

**DOI:** 10.64898/2026.02.19.706780

**Authors:** Ioannis Grammatikakis, Chosita Norkaew, You Jin Song, Amit K. Behera, Erica C. Pehrsson, Corrine Corrina R. Hartford, Shreya Kordale, Rishabh Prasanth, Yongmei Zhao, Biraj Shrethsa, Xiao Ling Li, Ravi Kumar, Ragini Singh, Tayvia Brownmiller, Xinyu Wen, Natasha J. Caplen, Pablo Perez-Pinera, Kannanganattu V. Prasanth, Thomas Gonatopoulos-Pournatzis, Ashish Lal

## Abstract

Intron retention (IR) is increasingly recognized as a feature of long noncoding RNAs (lncRNAs), yet the mechanisms that shape IR in lncRNAs and the functional consequences of this process remain largely unexplored. To investigate how IR contributes to lncRNA regulation, we performed a genome-wide screen to identify factors controlling IR in the lncRNA *PURPL*. This approach uncovered a prominent role for U2AF2, which promotes retention of a specific intron in *PURPL* through a weak polypyrimidine tract. IR of this intron drives nuclear enrichment of *PURPL* and enhances cell proliferation, revealing biological relevance. Transcriptome-wide analyses showed that although U2AF2 broadly supports canonical splicing consistent with its well-established function in promoting splicing, it also facilitates IR within a distinct subset of RNAs, including the nuclear speckle–associated lncRNA *MALAT1*. Loss of U2AF2 disrupts *MALAT1* speckle localization and using *MALAT1* knockout cells reconstituted with wild-type or intron deleted variants, we identified a single intron critical for *MALAT1*’s speckle localization. Deletion of this intron from endogenous *MALAT1* impaired speckle localization and reduced cell migration, phenocopying the loss of *MALAT1*. Together, these findings reveal IR as a key regulatory mechanism governing lncRNA localization and function and uncover an unexpected role for U2AF2 in promoting IR within specific lncRNA contexts.

## Introduction

Long non-coding RNAs (lncRNAs) are RNA molecules >200 nucleotides (nt) long that lack coding potential. Similar to protein-coding genes, lncRNAs are transcribed by RNA Polymerase II, are 5′-capped, spliced and are often polyadenylated (1). Several characteristics distinguish lncRNAs from protein-coding genes, including higher tissue-specific expression, lower abundance and stability, predominant nuclear localization, and having fewer but longer exons (2–6). One of the most distinguished features of lncRNAs is that they are less efficiently spliced compared to mRNAs (4,7–9). Interestingly, lncRNAs contain introns with much slower intron-excision kinetics compared to protein-coding genes (10). However, the factors that regulate lncRNA splicing and their impact on lncRNA biology remain largely unclear. We previously reported that the lncRNA *PURPL* exhibits prominent splicing defects, including frequent retention of its second intron. Transcriptomic analyses using both long-read and short-read sequencing revealed multiple *PURPL* isoforms, several of which retain intron 2 (11). While *PURPL* is also known to be transcriptionally induced by p53 and to contribute to cancer cell growth through multiple mechanisms, the regulatory factors governing intron 2 splicing and the functional significance of its retention have not been defined. This gap provides an opportunity to uncover broader principles of IR regulation in lncRNAs.

Splicing is a highly regulated process where *cis*- and *trans*-acting factors coordinate proper and efficient spliceosomal recruitment (12). Several consensus sequences in the RNA are crucial for splicing such as the polypyrimidine (Py)-tract, a stretch of pyrimidines close to the 3′ splice site (ss) (13). The splicing factor U2AF1 (U2-auxiliary factor 1) binds to the 3′ ss and U2AF2 binds to the Py-tract. The U2AF heterodimer is recruited to the branch point, and this complex serves as a critical intermediate for the subsequent assembly of the spliceosome and splicing catalysis (14–18). The vast majority of human genes encode pre-mRNAs that contain introns that are spliced out and most nascent transcripts undergo alternative splicing, which leads to the combinatorial joining of exons. Alternative splicing is a major contributor to protein isoform diversity, enabling the generation of multiple transcript variants from a single gene and frequently shaping measurable cellular phenotypes (19–22). One of the least studied forms of alternative splicing is intron retention (IR). Lately, it has drawn more attention since it has an important role in post-transcriptional gene expression regulation, and it is believed that more than half of human multi-exonic genes are affected by IR (23). IR frequently leads to nuclear detention of transcripts and eventually degradation which may serve as a defense mechanism of the cell against deleterious products. The extra sequence added by the retained intron may change the open reading frame (ORF) of protein-coding genes and lead to nonsense-mediated decay (NMD) (24–26) or initiate the use of an upstream ORF (27,28). In addition, they can contain binding sites for microRNAs (miRNAs) or RNA-binding proteins (RBPs), or they may form structural changes that impact the function of the transcript (29). Interestingly, IR has been associated with several diseases such as cancer where tumor cells show higher levels of IR compared to normal tissues (30). Studies on IR have shown that retained introns have been associated with weaker (non-consensus) splice sites, higher GC content, various lengths of introns, and higher intronic sequence conservation (23,31). lncRNAs have been shown to participate in splicing regulation, with *MALAT1* as the most prominent example. *MALAT1* is a highly abundant nuclear lncRNA which has oncogenic activity and is one of the first lncRNAs associated with cancer (32–34). It also has the ability to regulate alternative splicing because it resides in the nuclear speckles and can interact with several SR (Serine/Arginine rich) proteins to regulate their phosphorylation state and distribution between transcription sites and speckles (33,35–37). Nuclear speckles are sub-nuclear compartments that act as hubs in coordinating transcription, RNA processing and mRNA export (38). Several RBPs reside in nuclear speckles such as the protein SON which is a key component and marker of nuclear speckles (39). However, *MALAT1* itself is not essential for the structural integrity of nuclear speckles (40,41). Nevertheless, it remains unclear whether *MALAT1* undergoes alternative splicing and how this process impacts its localization to nuclear speckles.

Here, we show that the intron 2-retaining *PURPL* isoform, which localizes to the nucleus, has a shorter half-life than the spliced isoform and promotes cell proliferation. To identify regulatory factors controlling the splicing of this functionally relevant intron, we utilized our recently developed CRISPR-based genome-wide screening strategy using a pooled guide RNA library with a *PURPL* intron 2 minigene reporter containing its flanking exons. Unexpectedly, this approach revealed a role of the splicing activator U2AF2 in promoting *PURPL* IR. U2AF2 was further found to promote a broader intron retention program that includes two introns within *MALAT1*. Notably, depletion of U2AF2 resulted in *MALAT1* exclusion from nuclear speckles. *MALAT1* knockout cells exogenously expressing *MALAT1* truncation mutants lacking specific introns identified one intron essential for the nuclear speckle association of the *MALAT1* transcript. Breast cancer cells where this intron is deleted showed reduced cell migration demonstrating a functional role of the intronic sequences. These findings provide insights into the splicing regulation of *PURPL* and *MALAT1*, uncovering a previously unrecognized functional role of intron retention in lncRNAs.

## Results

### A CRISPR-based screen identifies U2AF2 as an activator of *PURPL* intron 2 retention

We previously identified *PURPL* as a p53-regulated lncRNA that promotes the growth of colorectal and liver cancer cells (11,42). In those studies, we reported that *PURPL* undergoes alternative splicing, with the ∼1.9 kb intron 2 either retained or spliced in the mature transcript (11). To identify regulators of *PURPL* intron 2 retention, we applied our recently developed screening platform, CRISPR-based identification of Regulators of Alternative Splicing with Phenotypic Sequencing (CRASP-seq) (43). This approach utilizes a genome-wide knockout lentiviral library targeting ∼18,500 protein-coding genes using a dual hybrid guide (hg)RNA system that combines *Streptococcus pyogenes* (*Sp*)Cas9 and optimized *Acidaminococcus sp.* (op)Cas12a nucleases. (22,44–46). The library also includes ∼2,400 pairs and ∼500 curated genetic interaction pairs, allowing combinatorial perturbations from a single construct. Each gene or gene pair is targeted by four independent hgRNAs alongside ∼500 intergenic and non-targeting controls, totaling ∼ 91,000 hgRNAs. The lentiviral library incorporates a doxycycline-inducible *PURPL* IR reporter comprised of the full native intron 2 sequences (∼1.9 kb) flanked by the adjacent exons 2 (161 bp) and 3 (163 bp) (Figure 1A).

**Figure 1.**
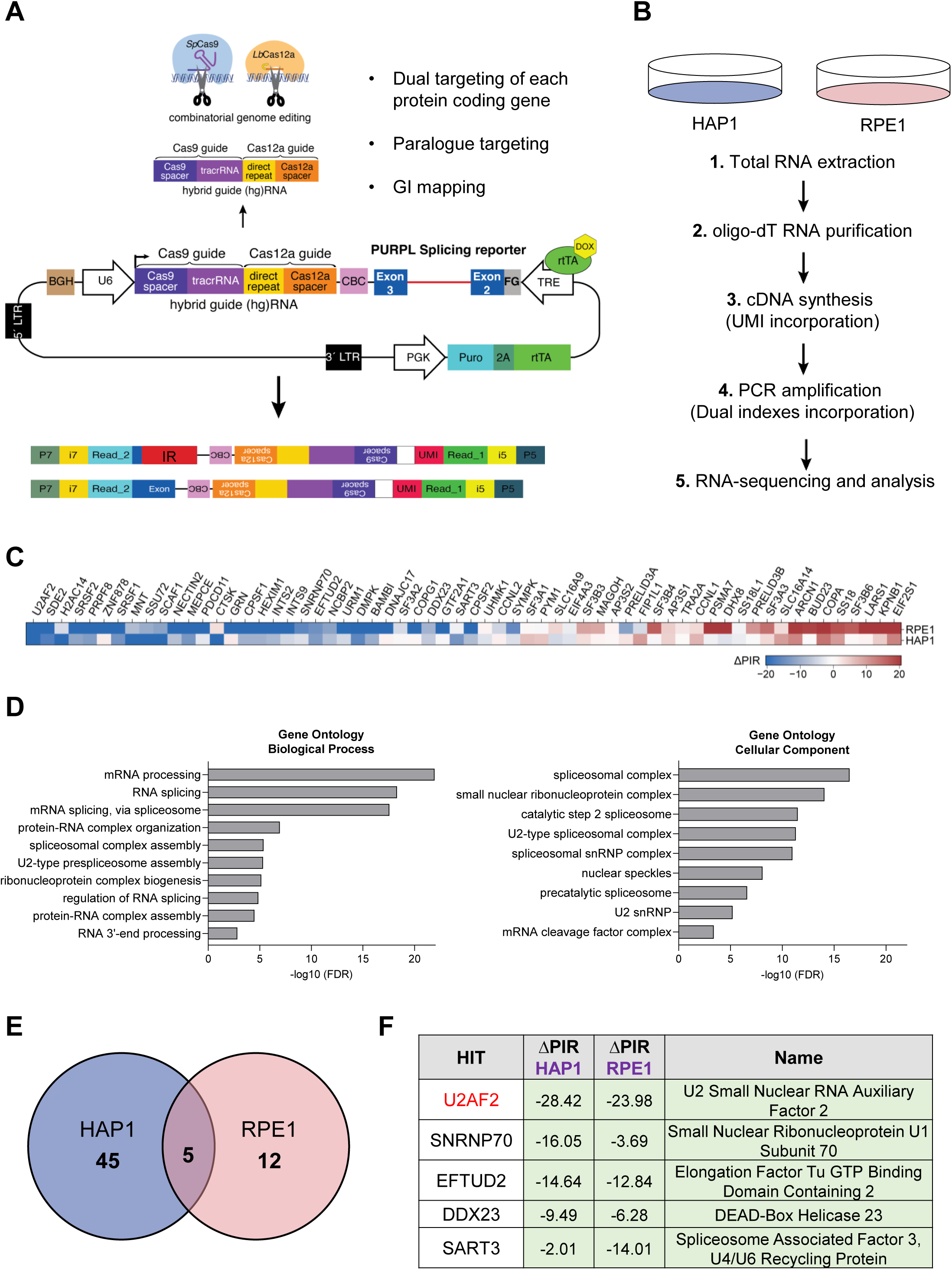
CRISPR-based screen identifies U2AF2 as a promoter of intron retention. **(A)** Schematic of the *PURPL* IR reporter used in the screen to identify IR regulators. The pLCHKO vector contains the puromycin resistance gene used for selection. The transgene also contains the genomic sequences of exon 2 (161 bp), intron 2 (1,943 bp), and exon 3 (163 bp) of the *PURPL* gene under a Doxycycline-inducible promoter. A hybrid guide (hg)RNA for guiding *Sp*Cas9 and *Lb*Cas12a is expressed under a U6 promoter. Top arrow: The hgRNA is cleaved by Cas12a to produce 2 guide RNAs for dual targeting of each protein coding gene, paralog targeting, and genetic interaction mapping. Bottom arrow: Upon intron splicing or intron retention, two different reads are generated. **(B)** Two different cell lines, stably expressing Cas9 and Cas12a, are used: HAP1, and RPE1. The CRASP-seq pipeline until Illumina paired-end sequencing of CRISPR libraries is depicted. **(C)** Heatmap showing single genes only (after removing gene pairs), whether identified individually or as part of a gene pair. The hits are ordered by the average ΔPIR values across HAP1 and RPE1, from lowest to highest. **(D)** Left: Gene ontology enrichment analysis of biological processes regulated by the hits of the screen. The most significant pathways involve mRNA processing and RNA splicing. Right: Gene ontology enrichment analysis of cellular components regulated by the hits of the screen. These components include mainly Spliceosomal Complex, small nuclear ribonucleoprotein complex and the Catalytic step 2 Spliceosome. **(E)** Venn diagram showing that there are 5 common hits which cause a reduction in the percentage of intron retention (ΔPIR) of the splicing reporter in the two cell lines. **(F)** Table showing the top 5 hits of the screen with a ΔPIR for each hit in each cell line and their full names.

We performed the CRASP-screen in two cell lines: HAP1 and RPE1. Following puromycin selection for 48 h and doxycycline induction for an additional 24 h, RNA was extracted, and poly-A RNA was purified using oligo(dT) beads to exclude transcripts still undergoing processing. cDNA was synthesized using customized primers containing unique molecular identifiers (UMIs), facilitating the removal of PCR duplicates during downstream analysis. The synthesized cDNA was then utilized to construct Illumina sequencing libraries for paired-end sequencing. This sequencing approach enables simultaneous quantification of transcripts that either retain or splice out an intron of interest from one read, while the corresponding hgRNA sequences were captured in the other read to pinpoint specific genetic perturbations. By directly linking *PURPL* splicing outcomes with genetic modifications introduced by each hgRNA in the library, this platform allowed the genome-wide identification of *PURPL* splicing regulators (Figure 1B).

We employed an analytical pipeline that prioritized candidate genes based on multiple hgRNAs showing significant deviations from intergenic controls, with consistent identification across two technical replicates. This analysis identified 62 genes that influence *PURPL* intron 2 splicing, the majority of which act as suppressors of intron 2 splicing (Figure 1C and Table S1). Gene ontology analysis of biological processes revealed a strong enrichment for terms related to mRNA processing and RNA splicing. Additionally, analysis of cellular components indicated that the identified hits are predominantly associated with Spliceosomal Complex, including the U2-type Spliceosomal Complex and the Catalytic step 2 Spliceosome (Figure 1D). These findings validate the effectiveness of our screening approach.

Consistent with the endogenous *PURPL* transcript isoforms, the basal levels of IR of the reporter were very high, with the majority of amplified transcripts corresponding to intron-containing isoforms, as expected based on the intensity of the Illumina libraries before sequencing (Figure S1A). Thus, the detection of genes that suppress IR might be below the detection levels of this screen. Therefore, we focused on the genes that promote intron retention. By calculating the change in Percent Intron Retention (ΔPIR = PIR_guide_ – PIR_intergenic_) across the two cell lines, we identified 5 unique genes as regulators of *PURPL* intron 2 retention, all of which act to suppress intron 2 splicing (Figure 1E and Table S1). This highly significant overlap between the two cell lines (Figure 1E; odds ratio = 204.4, p=2.81xe^-10^; Fisher’s exact test) indicated a shared regulatory mechanism governing *PURPL* IR, although cell-type-specific effects may also contribute.

Surprisingly, the gene with the strongest impact based on the average ΔPIR metric and identified as a hit in both cell lines, was U2AF2 (Figure 1F). This was followed by Small Nuclear Ribonucleoprotein U1 Subunit 70 (SNRNP70), a core component of the U1 snRNP, and Elongation Factor Tu GTP Binding Domain, which is associated with U5 snRNP (Figure 1F). Our finding that U2AF2 promotes IR of *PURPL* intron 2 was unexpected because U2AF2 is a component of the spliceosome machinery, and its canonical function is to promote intron removal (14,18).

### U2AF2 binds to the *PURPL* intron 2 transcript and promotes intron retention

U2AF2, as a heterodimer with U2AF1, binds to the Py-tract and the 3′ ss, respectively, to promote splicing (14,17). To elucidate its role in *PURPL* splicing, we investigated whether U2AF2 binds to intron 2. Enhanced Crosslinking and Immunoprecipitation sequencing (eCLIP-seq) (47), from the ENCODE project (48,49) from HepG2 cells revealed multiple U2AF2 and U2AF1 binding sites not only proximal to the 3′ ss of *PURPL* intron 2 but also broadly distributed across intron 2 (Figure 2A). Compared to eCLIP data for other RBPs, U2AF1 and U2AF2 showed the highest density of binding sites and the most prominent peaks. Specific binding of U2AF2 to intron 2 was confirmed by performing RNA immunoprecipitation (RNA-IP) using a U2AF2-specific antibody and primer pairs designed to detect *PURPL* transcripts. U2AF2 was found to interact with transcripts containing *PURPL* intron 2, but not with the spliced *PURPL* transcripts (Figure 2B and S1B).

**Figure 2.**
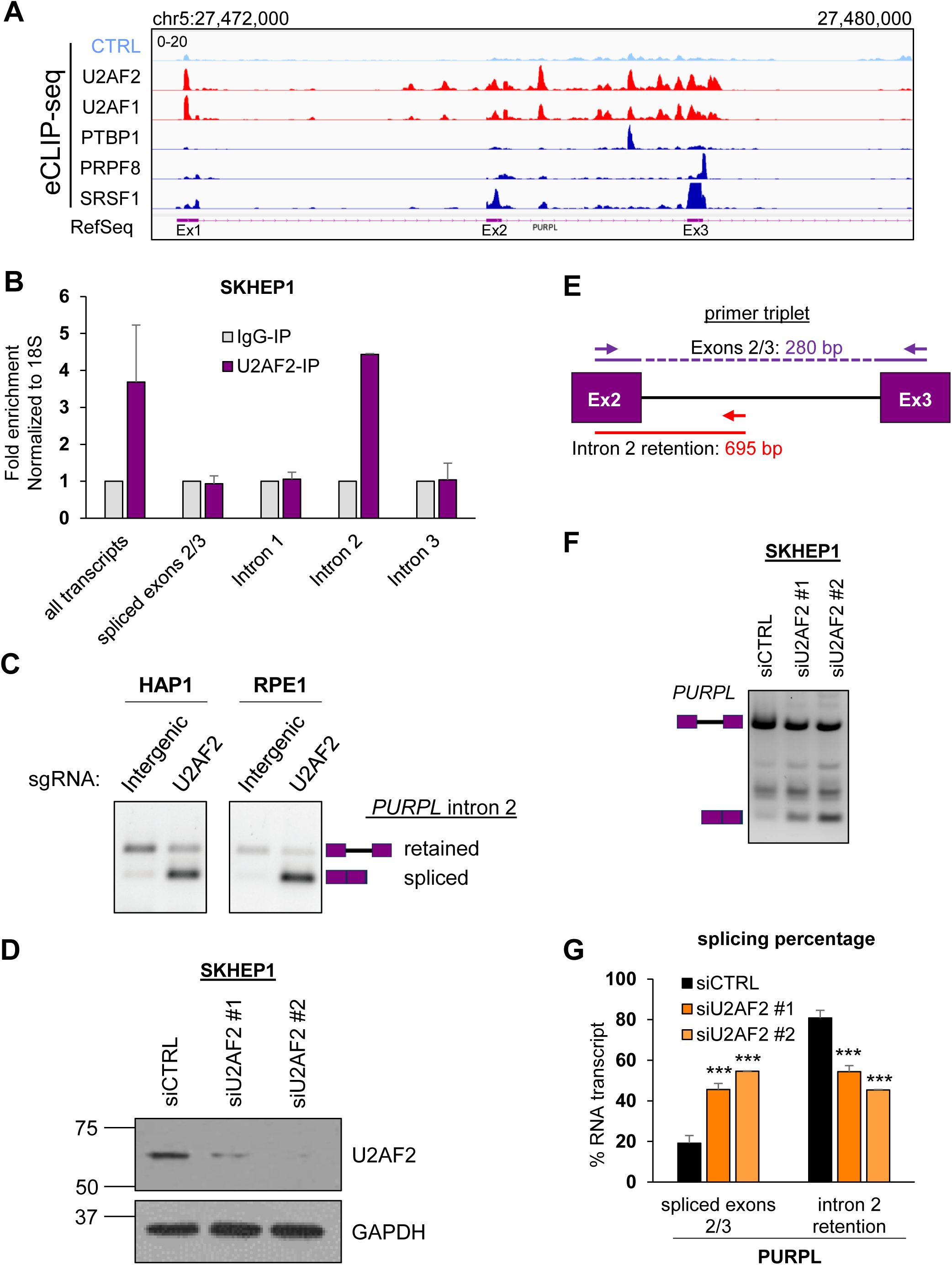
U2AF2 binds to intron 2-containing *PURPL* transcripts and promotes intron 2 retention. **(A)** IGV snapshot of eCLIP-seq data showing binding sites and enrichment of U2AF2 and U2AF1 on the *PURPL* transcripts around intron 2. eCLIP-seq data for PTBP1, PRPF8, and SRSF1 are indicated. The annotated locus by RefSeq is also indicated. eCLIP data-seq was downloaded from encodeproject.org. **(B)** RT-qPCR after RNA-IPs using a U2AF2 antibody with primer pairs specifically detecting *PURPL* transcripts as indicated in Figure S1B. U2AF2 binds to transcripts containing intron 2 but not the ones with intron 1, intron 3, or spliced exons 2 and 3. Samples were normalized to IgG-IP. 18S was used as a loading control. **(C)** Validation of hgRNAs using the *PURPL* minigene in HAP1 and RPE1 cells. The gels show RT-PCR products for *PURPL* transcripts with specific primers for the minigene. The hgRNAs used were targeting intergenic region, or U2AF2. The schematics next to the gel indicate the expected products of the intron-retained and spliced isoforms. **(D)** Western blot for U2AF2 showing successful knockdown of U2AF2 protein in SKHEP1 cells. GAPDH was used as a loading control (lower panel). **(E)** Schematic of the PCR primer triplet used to detect intron 2 retention (red) or splicing (purple). The length for each PCR product is indicated. **(F)** Gel with RT-PCR products for *PURPL* upon knockdown of U2AF2 with 2 different siRNAs in SKHEP1 cells. The schematics next to the gel indicate the expected products of the intron-retained and spliced isoforms. Between the two expected PCR products, we observed an extra band corresponding to the inclusion of an alternative exon inside *PURPL* intron 2 as observed in RefSeq, the inclusion of which is not affected by U2AF2. **(G)** Graph with quantitation of the gel bands in **(F)**. Error bars represent standard deviations from 3 **(G)** experiments. ***p<0.001.

To functionally validate the role of U2AF2 in *PURPL* intron 2 retention, we performed RT-PCR assays from HAP1 and RPE1 cells transduced with *PURPL* minigene reporter constructs and control or *U2AF2*-targeting sgRNAs. In both cell lines, we observed a striking decrease in intron 2 retention upon *U2AF2* depletion as compared to control cells expressing sgRNAs targeting intergenic regions (Figure 2C). As an alternative approach, we knocked down *U2AF2* using two independent siRNAs and conducted semi-quantitative RT-PCR assays from SKHEP1, HepG2, and normal human diploid fibroblasts W138 cells, using a primer triplet (Figures 2D-2F; Figures S2A and S2B). In all tested cell lines, U2AF2 depletion increased *PURPL* intron 2 splicing, from ∼20% to ∼40-60%, more than doubling the abundance of the spliced isoform. Given that U2AF2 functions as a heterodimer with U2AF1, we also knocked down *U2AF1* in SKHEP1 cells and observed an effect similar to *U2AF2* knockdown on *PURPL* intron 2 retention regulation (Figure S2C). We expected that U2AF1 would also come as one of the top hits in the screen but *U2AF1* sgRNAs were not included in the library due to high off-target scores of the guides. These results demonstrate that the U2AF complex promotes *PURPL* intron 2 retention and that both U2AF1 and U2AF2 promote intron retention in *PURPL*.

### The intron-retained *PURPL* transcript is unstable and localizes to the nucleus

We previously observed an enrichment of the intron 2-retaining *PURPL* transcript in the nucleus in liver cancer cells (11). To validate these findings, we conducted single molecule RNA-FISH in HCT116 cells using probes hybridizing to *PURPL* intron 2 (Figure 3A). Because *PURPL* is upregulated in response to DNA damage (42), we treated the cells with hydroxyurea for 24 h to induce *PURPL* expression. RNA-FISH revealed that the *PURPL* intron 2 containing transcript was exclusively localized to the nucleus. To assay for subnuclear compartmentalization, we also performed RNA-FISH to detect *MALAT1*, which resides in nuclear speckles (36). Intron 2-retained *PURPL* transcripts did not colocalize with *MALAT1* indicating that *PURPL* is excluded from nuclear speckles (Figure 3A). We further validated this result by conducting nuclear/cytoplasmic fractionation assays followed by RT-qPCR and found that *PURPL* intron 2-containing transcripts were almost exclusively nuclear (Figure S3A).

**Figure 3.**
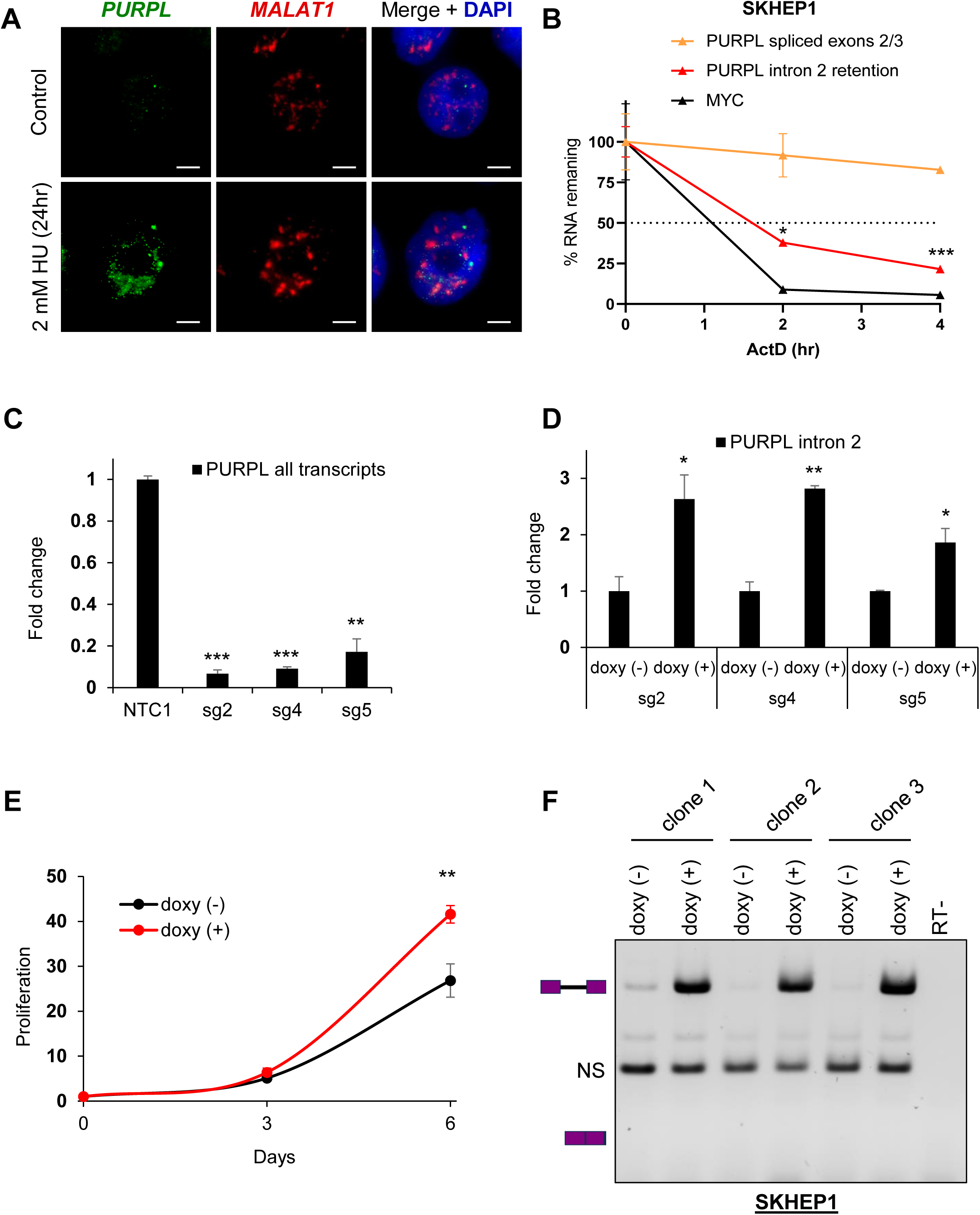
Expression of Intron 2 Retaining *PURPL* leads to high proliferation. **(A)** RNA-FISH images for *PURPL* with intron 2 retention and *MALAT1* in HCT116 cells without treatment or after 24 h of 2 mM of Hydorxyurea (HU) to induce *PURPL* expression. Scale bar is 10 μm. **(B)** RNA stability assays were performed for *PURPL* transcripts by measuring their levels by RT-qPCR following 0, 2, and 4 h of ActD treatment in SKHEP1 cells. *MYC* was used as a positive control for unstable RNA. **(C)** RT-qPCR showing *PURPL* depletion in in SKHEP1 cells using 3 different gRNAs (sg2, sg4, sg5) compared to a Non-Targeting Control gRNA. **(D)** RT-qPCR for intron 2-containing PURPL transcript after 48 h of 1 μg/mL doxycycline treatment in comparison to no treatment in SKHEP1 *PURPL*-CRISPRi populations using the 3 gRNAs as in **(C)**. **(E)** Proliferation assay showing the effect of overexpression of intron 2-containing *PURPL* transcript in the proliferation of SKHEP1 cells where the endogenous *PURPL* is knocked down with CRISPRi. Error bars represent standard deviation from 3 populations with different gRNAs. The cells were treated with 1 μg/mL doxycycline to induce intron 2-retained *PURPL* expression and cell proliferation was monitored at 3 and 6 days. **(F)** Gel with RT-PCR products for transcripts using primers as indicated in Fig. 2E upon doxycycline treatment of SKHEP1 CRIPSRi cells. The three repeats represent 3 different clones of cells. The last lane is an RT- control. The schematics next to the gel indicate the expected products of the intron-retained and spliced isoforms. NS indicates a non-specific band. Error bars in **(B)**, **(C)**, and **(D)** represent standard deviations from 2 experiments. *p<0.05, **p<0.01, ***p<0.001.

Intron retention can affect transcript stability, with longer RNA species often being more vulnerable to degradation. To test if *PURPL* intron 2 effects transcript turnover, we conducted RNA stability assays using the transcription inhibitor Actinomycin D (ActD) and observed that the intron 2-retained *PURPL* transcript was less stable and had a significantly lower half-life than the *PURPL* transcript in which intron 2 was spliced out (Figure 3B). To determine the functional impact of *PURPL* intron 2 retention, we used CRISPRi to generate *PURPL*-depleted SKHEP1 cells by targeting 3 independent sgRNAs to the p53-response element of the *PURPL* promoter that we had used in our previous study (11). We observed >80% reduction in the levels of endogenous *PURPL* in these experiments (Figure 3C). We next stably transduced these *PURPL*-depleted cells with a lentivirus to express exogenous *PURPL* transcript specifically containing intron 2 under the control of a doxycycline (doxy)-inducible promoter and observed ∼2-3-fold induction of the exogenous *PURPL* intron 2 containing transcript (Figure 3D). Because we and others have previously shown that *PURPL* is overexpressed in some cancers and *PURPL* promotes proliferation (11,42,50,51), we next performed cell proliferation assays from the doxy-inducible cells. Our results showed that cells expressing intron 2-contaning *PURPL* proliferate faster (∼2-fold) as compared to the no-doxy cells after 6 days of doxy treatment (Figure 2E). Induction of intron 2-contaning *PURPL* transcripts did not have any effect on live cell percentage which remained at ∼100% (Figure S3B). Importantly, as a negative control, there was no effect of doxy treatment on the proliferation of parental SKHEP1 cells (Figure S3C). To confirm that proliferation increase induced by exogenous *PURPL* transcript containing intron 2 was not due to the spliced isoform, we performed semi-quantitative RT-PCR. As shown in Figure 3F, doxy treatment resulted exclusively in the detection of the intron-retained *PURPL* transcript. These data suggest that intron 2 retention modulates *PURPL* function by sequestering the transcript in the nucleus and reducing its stability, while simultaneously promoting cell proliferation, highlighting a critical regulatory role for intron 2.

### A weak polypyrimidine tract is critical for *PURPL* intron 2 retention

U2AF2 binds to the Py-tract to initiate spliceosomal assembly leading to splicing and intron removal (14). Suboptimal splicing efficiency may be the result of multiple factors, including splice site strength and the Py-tract sequence composition. The Py-tract is a stretch of 15-20 pyrimidines (mostly Us) immediately upstream of the 3′ ss. We analyzed the Py-tract of *PURPL* intron 2 and noticed that it deviates from the consensus, containing numerous purines (Figure S4A). To investigate whether U2AF2 regulation on intron 2 retention relies on the Py-tract sequence, we used the *PURPL* intron 2 minigene reporter system (Figure 1A). We mutated the Py-tract, replacing it either with a strong Py-tract from *PURPL* intron 1 or with a stretch of Us, simulating a strong Py-tract. *U2AF2* was knocked down in SKHEP1 cells stably expressing the mutated Py-tract constructs and RT-PCR was conducted to monitor intron 2 splicing. Interestingly, the basal levels of intron retention were very different with different mutants; ∼90% in intron 2 WT (wild-type), ∼30% in *PURPL* intron 1 and ∼20% in U-rich construct (Figure S4B and S4C). This data demonstrates that the weak Py-tract is responsible for the inefficient intron 2 removal in *PURPL* (Figure S4C). *U2AF2* knockdown resulted in modest increase in splicing of the mutant constructs as well. Because the minigene reporter contained the mutant Py-tract along with the remaining endogenous sequences from *PURPL* intron 2, the modest increase in splicing observed for the mutants after *U2AF2* knockdown may reflect U2AF2 inhibitory binding at additional sites within *PURPL* intron 2 beyond the Py-tract (Figure 2A). These data indicate that the endogenous intron 2 Py-tract in *PURPL* is a weak Py-tract that controls the high intron retention observed in *PURPL*.

### U2AF2 directly regulates multiple IR events including *MALAT1*

To determine if U2AF2 promotes IR in additional genes, we next performed total RNA-seq and PacBio Iso-seq from SKHEP1 cells transfected with siCTRL or siU2AF2. Consistent with our data, *U2AF2* knockdown resulted in a marked increase in the splicing efficiency of *PURPL* intron 2 (Figures S5A and S5B). Additionally, *U2AF2* depletion increased overall *PURPL* expression, with increased read coverage across all exons (Figures S5A and S5B; Tables S3 and S4). To identify IR events regulated by U2AF2 at a transcriptome-wide level, we utilized IRFinder, an algorithm specifically designed to calculate changes in IR (52). This analysis revealed ∼1000 statistically significant IR events regulated by U2AF2. For majority of these events (732), there was an increase in the IR ratio (Intronic abundance/(Intronic abundance + exonic abundance)) after *U2AF2* depletion, indicating that U2AF2 predominantly promotes splicing, as expected. However, in 269 events, *U2AF2* knockdown resulted in decreased IR, indicating that U2AF2 promotes IR in a subset of genes (Figure 4A and Table S2). We next analyzed ENCODE U2AF2 eCLIP-seq data from HepG2 cells to determine if U2AF2 directly binds to the regulated transcripts. U2AF2 binding was detected in 96 out of the 269 IR events supporting a direct role for U2AF2 in promoting IR (Figure S5C and Table S2).

**Figure 4.**
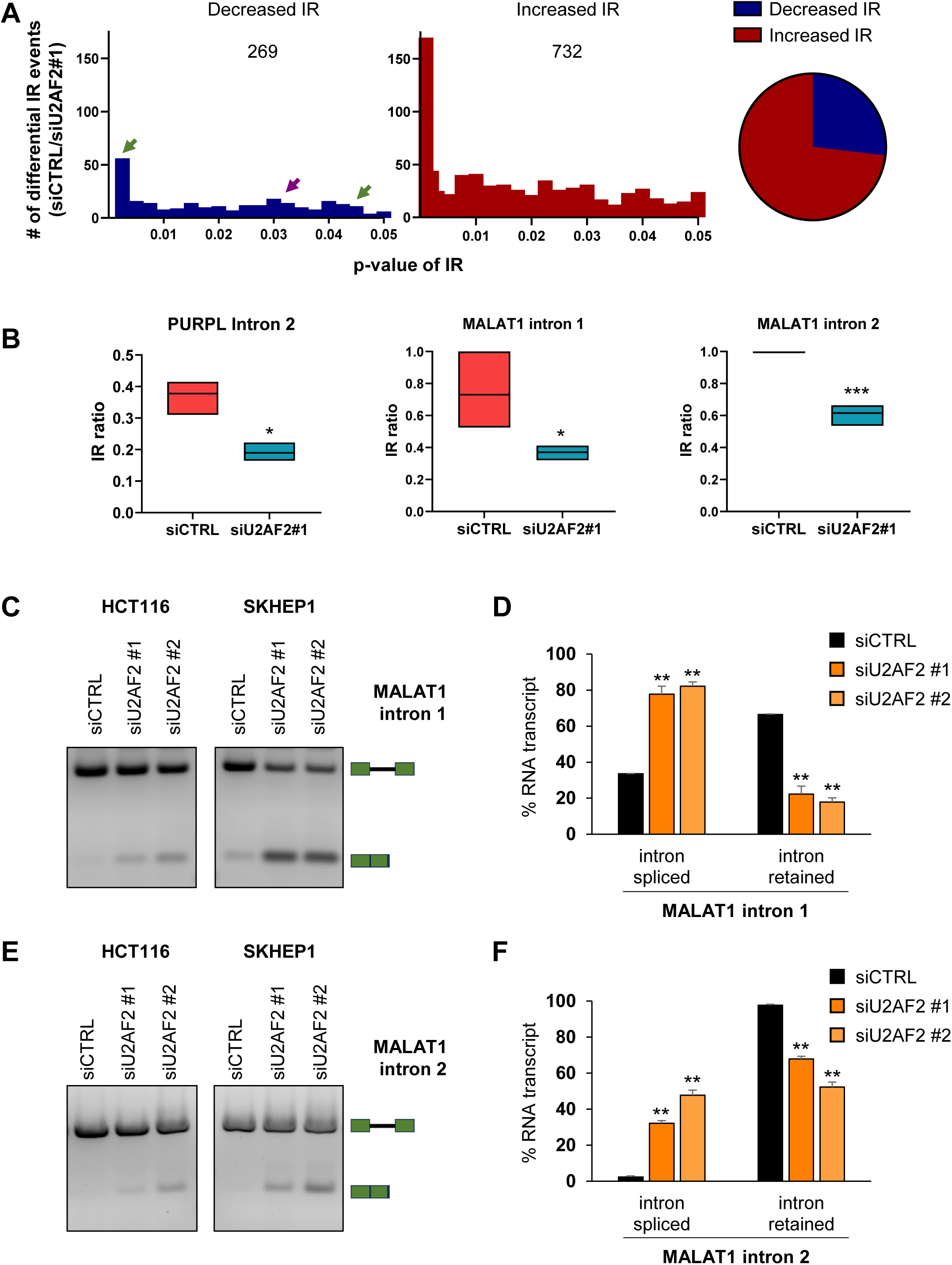
U2AF2 directly promotes Intron Retention of *MALAT1*. **(A)** U2AF2 was knocked down in SKHEP1 cells and 72 h later, RNA was extracted and RNA-seq was performed. *Left*: Number of decreased (blue) and increased (red) IR events at various p-values after U2AF2 knockdown as analyzed with the IR Finder algorithm. The purple arrow indicates the *PURPL* IR event and green arrows indicate *MALAT1* IR events *Right*: Pie chart of the numbers of increased and decreased IR events upon U2AF2 knockdown. **(B)** Floating bar plots showing the IR ratio of *PURPL* intron 2 and the IR ratio of intron 1 (middle) and intron 2 (right) of *MALAT1* in siCTRL and siU2AF2#1 samples as analyzed with the IRFinder algorithm. **(C)** and **(E)** RT-PCR for *MALAT1* using a primer pair flanking the regulated intron 1 **(C)** or intron 2 **(E)** upon knockdown of U2AF2 with 2 different siRNAs in HCT116 and SKHEP1 cells. The schematics next to the gel indicate the expected products of the intron-retained and spliced isoforms. **(D)** and **(F)** Bar graph with quantitation of the gel bands from **(C)** and **(E)** in SKHEP1 cells. Error bars represent standard deviations from 2 independent experiments. *p<0.05, **p<0.01, ***p<0.001.

To validate these findings, we selected three of the identified introns from *MDM1*, *RETREG2*, and *TMEM41B* (Figure S6A). We observed multiple U2AF2 binding peaks within the regulated introns from *MDM1* (Figure S6B). We next assessed their splicing in SKHEP1 cells following *U2AF2* knockdown using two independent siRNAs. In all three cases, *U2AF2* depletion significantly enhanced intron removal (Figures S6C-S6E). IRFinder validated that *PURPL* intron 2 was one of the events showing a significant decrease (∼2-fold) in IR ratio upon *U2AF2* depletion (Figure 4B).

Among the lncRNAs in which IR is promoted by U2AF2, was the lncRNA *MALAT1*. IRFinder identified two introns in *MALAT1* that have weak Py-tract sequences similar to *PURPL* intron 2 (Figure S7A-S7C). For both introns, *U2AF2* knockdown resulted in a significant decrease (∼2-fold) in the IR ratio compared to siCTRL (Figures 4A and 4B). This result was confirmed by RT-PCR from HCT116 and SKHEP1 cells upon *U2AF2* knockdown (Figures 4C-4F).

### Intron 2 of *MALAT1* drives its localization to nuclear speckles

*MALAT1* regulates splicing by interacting with splicing factors in nuclear speckles.(36) To determine the impact of IR on *MALAT1* localization to nuclear speckles, we performed RNA-FISH for *MALAT1* upon *U2AF2* depletion in SKHEP1 and HCT116 cells. Remarkably, *U2AF2* knockdown resulted in a significant decrease in the ratio of *MALAT1* signal intensity in speckles relative to nucleoplasm (Figures 5A and 5B; Figures S8A and S8B). U2AF2 depletion did not affect total *MALAT1* levels as observed in our RNA-seq (Table S3). This reduction in *MALAT1* levels in nuclear speckles did not compromise the structural integrity of nuclear speckles as indicated by immunostaining for the RNA-binding protein SON, a well-established nuclear speckle marker (39) (Figures 5A and S8A) and is consistent with previous reports showing that *MALAT1* is not essential for the integrity of nuclear speckles (40,41). These data suggest that U2AF2 plays an important role in promoting *MALAT1* localization to nuclear speckles.

**Figure 5.**
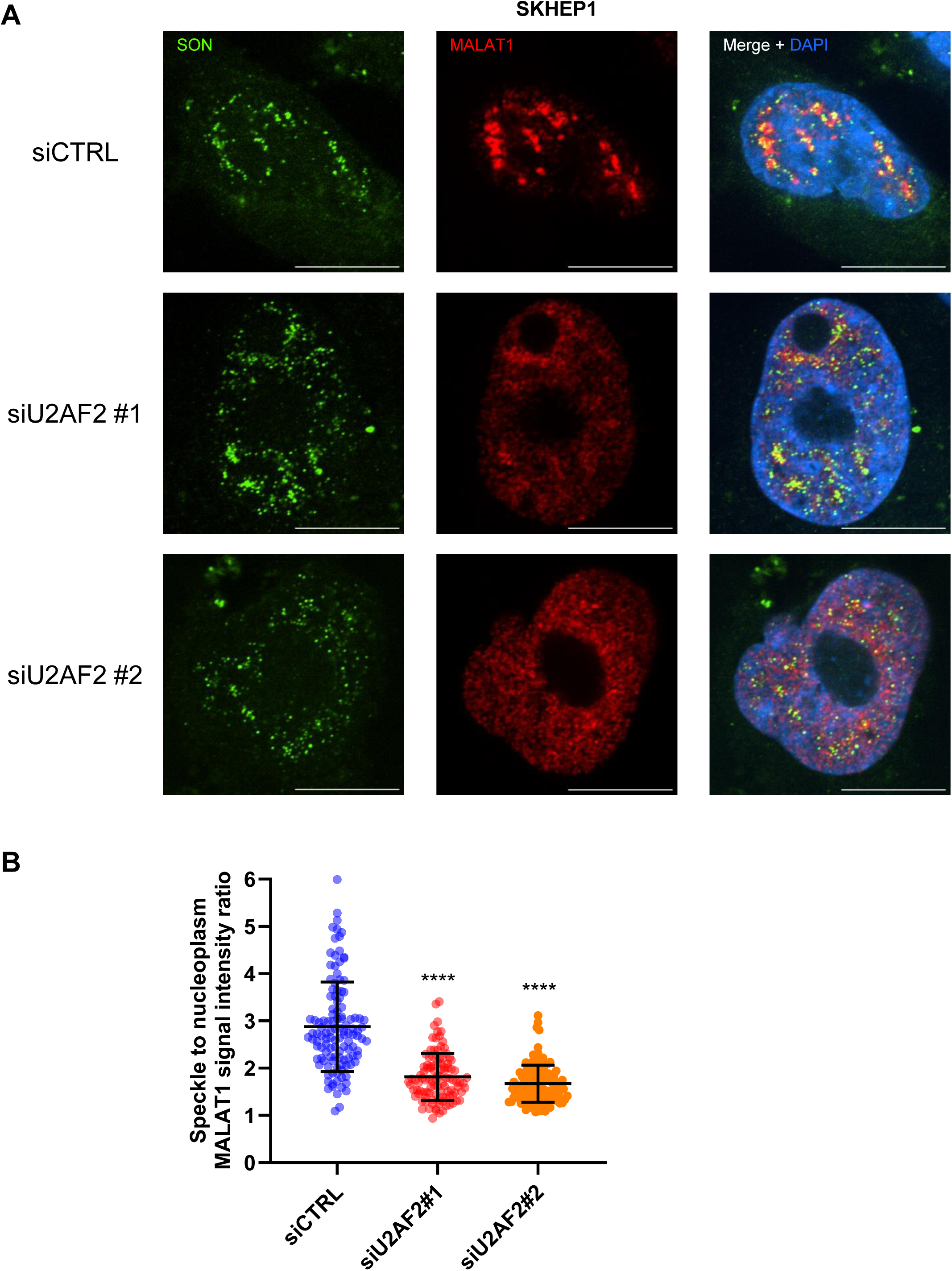
U2AF2 promotes localization of *MALAT1* to nuclear speckles. **(A)** RNA-FISH images for *MALAT1* and Immunofluorescence images for SON is shown upon transfection of SKHEP1 cells with siCTRL or siU2AF2 with two different siRNAs. *MALAT1* is enriched in nuclear speckles in the siCTRL but not upon U2AF2 knockdown. **(B)** Quantitation of the speckle to nuclear plasma *MALAT1* signal ratio in the three replicates in panel **(A)**. ****p<0.0001.

To determine whether it is the intron retention in the transcript *per se* which directs *MALAT1* localization or if U2AF2 indirectly regulates its localization, we assessed the nuclear speckle localization of wild-type (WT) and intron-deleted *MALAT1* transcripts. To do this, we used CRISPR/Cas9 to generate two *MALAT1* knockout (KO) clones in HCT116 by inserting ∼100 bp downstream of the transcription start site, a puromycin resistance gene in one allele and a hygromycin resistance gene in the other allele. In both KO clones, >95% reduction in *MALAT1* RNA levels was observed as compared to a wild-type clone (Figure S9A).

Next, we transduced the *MALAT1* KO clones with an empty vector or doxycycline-inducible lentiviral constructs that exogenously express full-length *MALAT1*-WT in which both introns are intact, intron 1 deleted *MALAT1* (del-1), intron 2 deleted *MALAT1* (del-2) or both introns were deleted (del-1+2). After 48 h of doxy-treatment to induce *MALAT1* expression, which was confirmed with RT-qPCR (Figure S9B), RNA-FISH combined with immunofluorescence for the nuclear speckle marker SON, revealed distinct localization patterns. As expected, exogenously expressed *MALAT1*-WT predominantly (∼80%) localized to nuclear speckles (Figures 6A and S10A). Similarly, *MALAT1*-del-1 also predominantly (∼80%) localized to nuclear speckles. In contrast, *MALAT1* lacking intron 2 (del-2) or both introns (del-1+2) exhibited a marked reduction (<15%) in nuclear speckle localization and instead displayed diffused localization in the nucleus (Figures 6A and S10A). Importantly, this diffused localization upon intron 2 deletion was observed in both *MALAT1* knockout clones and was further confirmed through quantitative analyses (Figures 6B and S10B).

**Figure 6.**
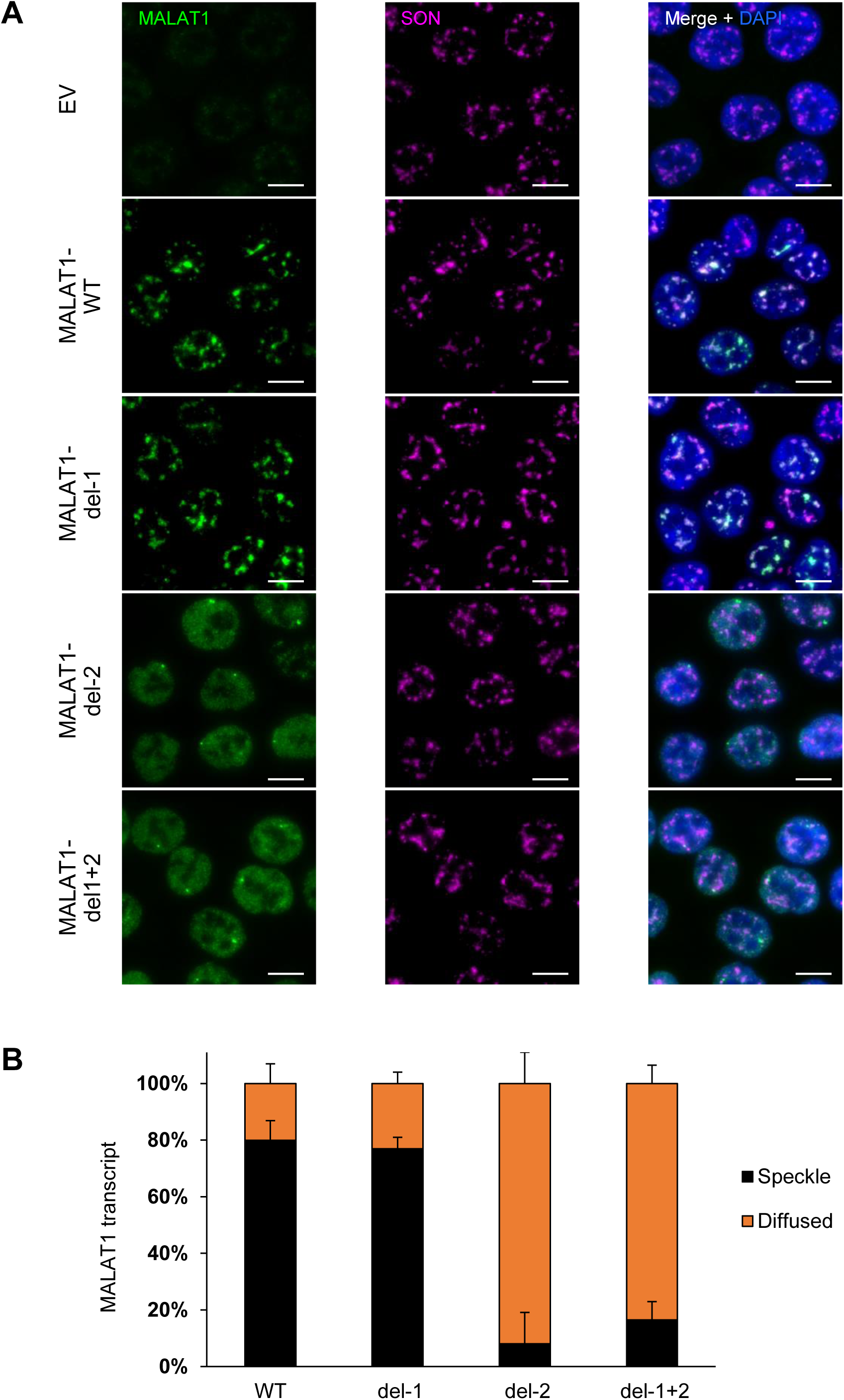
Intron 2 of *MALAT1* dictates its localization to nuclear speckles. **(A)** RNA-FISH images for *MALAT1* and Immunofluorescence images for SON in a HCT116 *MALAT1*-KO clone (clone 1) where Empty Vector (EV) or constructs that exogenously express *MALAT1* transcripts were re-introduced using a doxy-inducible lentivirus. Expressed *MALAT1* transcripts were full-length (WT) or intron 1 deleted (del-1) or intron 2 deleted (del-2) or both introns deleted simultaneously (del-1+2). The cells were first treated with 1 μg/mL doxycycline for 48 h to induce *MALAT1* expression. *MALAT1* without intron 2 does not enrich in speckles as it is shown with SON protein. Scale bar is 10 μm. **(B)** Graph showing quantification of the percentage of cells where *MALAT1* transcripts are enriched in speckles or being diffused in HCT116 cells clone 1. Error bars represent standard deviations from 2 independent experiments.

### Deletion of *MALAT1* Intron 2 using CRISPR/Cas9 impairs cell migration

*MALAT1* has been shown to promote cell migration and metastasis (32,53) and its expression has been correlated with high tumor progression and metastasis in various tumors (54). We therefore tested whether the intronic sequences that regulate *MALAT1* nuclear speckle localization are related to the ability of *MALAT1* to promote cell migration. To investigate this, we used the metastatic breast cancer cell line MDA-MB-231 and first performed RNA-FISH for *MALAT1* after knocking down *U2AF2*. Similar to SKHEP1 and HCT116 cells, *U2AF2* knockdown resulted in decreased localization of *MALAT1* to speckles in MDA-MB-231 cells (Figures 7A and 7B). We next utilized CRISPR/Cas9 to generate KO clones in MDA-MB-231 cells where either the whole *MALAT1* locus or intron 2 of *MALAT1* were deleted. *MALAT1* depletion in the *MALAT1* KO clone was confirmed with RT-qPCR (Figure S11A). RNA-FISH in 2 separate clones showed that intron 2 deletion reduced *MALAT1* localization to nuclear speckles compared to the WT clone (Figures 7C-7E). In the intron-2 deletion clones, the transcripts containing intron 2 were, as expected, undetected with primers specifically amplifying intron 2. Interestingly, there was a modest increase in total *MALAT1* expression suggesting a compensatory mechanism for the loss of intron 2 (Figure S11B). Functionally, when we tested the migration potential of these cells in transwell cell migration assays, we found significantly reduced cell migration in the intron 2-deleted clones, phenocopying the complete loss of *MALAT1* locus (Figures 7F and S11C). These data suggest that the retention of intron 2 is essential for *MALAT1* localization to nuclear speckles and it plays a key role in promoting cell migration. Together, our data uncovers a U2AF2-driven intron retention program in lncRNAs that regulates cell proliferation and migration.

**Figure 7.**
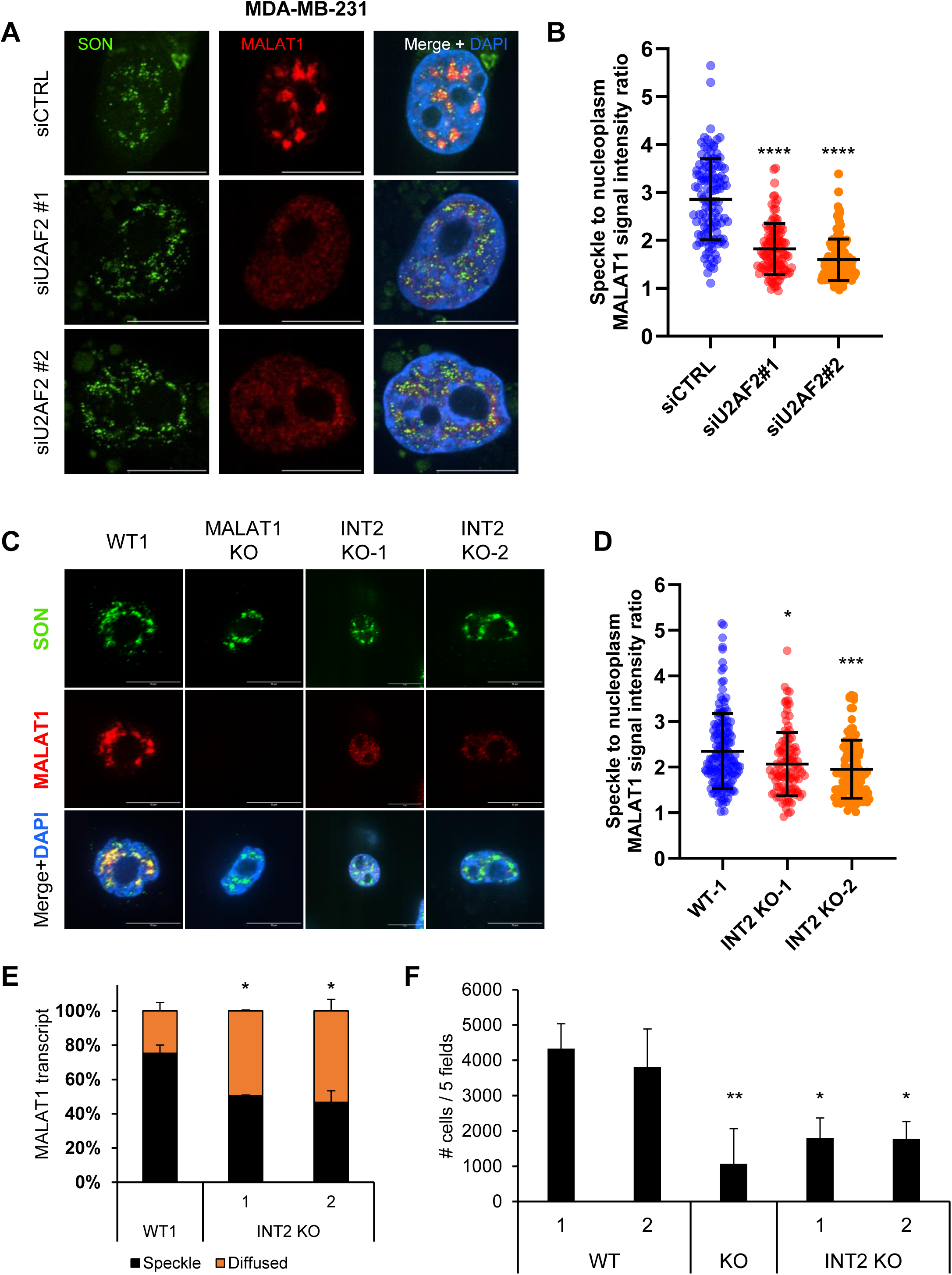
Intron 2 regulates the migration capacity of *MALAT1* in breast cancer cells. **(A)** RNA-FISH images for *MALAT1* and Immunofluorescence images for SON are shown upon transfection of MDA-MB-231 cells with siCTRL or siU2AF2. *MALAT1* is enriched in nuclear speckles in the siCTRL but not upon U2AF2 knockdown. **(B)** Quantitation of the speckle to nuclear plasma *MALAT1* signal ratio in the three replicates in panel **(A)**. **(C)** RNA-FISH images for *MALAT1* and Immunofluorescence images for SON in MDA-MB-231 clones either WT, or whole locus *MALAT1* deletion, or intron 2 deletion. When intron 2 is deleted, *MALAT1* does not enrich in speckles as it is shown with SON protein. Scale bar is 10 μm. **(D)** Quantitation of the speckle to nuclear plasma *MALAT1* signal ratio in the three replicates in panel **(C)**. **(E)** Graph showing quantification of the percentage of *MALAT1* transcripts enriched in speckles or being diffused in MDA-MB-231 cells in panel **(C)**, N=2. **(F)** Bar graph showing the number of cells that have migrated in transwell migration assays conducted in MDA-MB-231 WT, *MALAT1* KO and *MALAT1* – intron 2 deletion clones. Intron 2 deletion leads to decreased migration potential. The number of cells are the sum of cells from 5 different fields, N=4. *p<0.05, **p<0.01, ***p<0.001, ****p<0.0001.

## Discussion

The goal of this study was to examine the causes and consequences of IR in lncRNAs, focusing on *PURPL* and *MALAT1.* Intron retention is a regulated process that shapes gene expression networks through mechanisms such as NMD and influences alternative protein isoform generation during differentiation, development, and disease (25,55). We previously identified a p53-induced lncRNA, *PURPL*, which exhibits growth-promoting functions, and that intron 2 of *PURPL* is retained in multiple cell lines (11). To accomplish our goal, we employed an innovative RNA-seq coupled genome-wide CRISPR screen on a reporter containing *PURPL* intron 2 and unexpectedly identified U2AF2 as a major inhibitor of intron 2 removal in *PURPL*. U2AF2 forms a heterodimer with U2AF1, binding to the Py-tract and the 3′ ss, respectively, to promote splicing by recruiting core spliceosomal components (17,18,56). Therefore, given the established role of U2AF2 as a critical positive regulator of splicing catalysis, we were surprised that U2AF2 inhibits the removal of *PURPL* intron 2. While U2AF1 was not identified as a hit in our screen, follow-up experiments revealed that U2AF1 also suppresses *PURPL* intron 2 removal, suggesting a previously unrecognized role for the U2AF complex as a negative regulator *PURPL* IR.

Beyond IR in *PURPL*, we sought to determine whether U2AF2 promotes IR in other transcripts. Using IR Finder (52), we identified other genes where U2AF2 promoted IR. One of the genes that drew our attention was the lncRNA *MALAT1* which showed two intron retention events regulated and bound by U2AF2. These introns contain weak splice sites suggesting that this may be the reason that they are regulated, and they are not constitutively spliced. Because *MALAT1* interacts with splicing factors within nuclear speckles (36), we examined whether U2AF2 affects *MALAT1* localization to speckles. Indeed, our results establish *MALAT1* intron 2 as a critical determinant of its nuclear speckle localization. Future studies involving mapping of the intronic sequences and targeted mutagenesis of the splice sites within *MALAT1* intron 2 could help distinguish whether speckle localization of *MALAT1* is inhibited by the splicing process itself or depends on a discrete element within the intron.

For *PURPL*, previous work established its context-dependent oncogenic activity (11,42), but those studies depleted all transcripts from the *PURPL* locus despite the presence of multiple alternatively spliced isoforms. In the current study, overexpression of the intron 2-retaining *PURPL* isoform in CRISPRi-mediated *PURPL*-depleted cells increased proliferation, demonstrating isoform-specific, growth-promoting function. Our findings reveal that intron retention in both *MALAT1* and *PURPL* is a regulatory mechanism with direct consequences on lncRNA localization and cancer-related phenotypes.

The surprising finding of U2AF2 promoted IR promoted us to determine the underlying mechanism. We first asked whether U2AF2 influences transcription initiation or elongation at the *PURPL* locus, given the tight coupling between transcription and splicing (57,58). Transcription kinetics can modulate IR (23,59) and lncRNAs often lack proximal RNA-pol II phosphorylation over 5′ ss (10). We therefore assayed for PolII-pSer5 (transcription initiation) and PolII-pSer2 (transcription elongation) occupancy after U2AF2 knockdown using CUT&RUN-Seq but there was no significant difference around intron 2 or elsewhere across the *PURPL* locus (data not shown). We also examined the chromatin mark H3K36me3 which has been linked to IR (60) and again observed no effect after U2AF2 depletion (data not shown).

U2AF2 functions by binding to the Py-tract, and the Py-tract of *PURPL* intron 2 and *MALAT1* introns 1 and 2 deviate from the consensus, containing a mix of pyrimidines and purines. These atypical sequences likely contribute to splicing inhibition as revealed by our data from the mini-gene reporter for *PURPL*. Given that U2AF1 and U2AF2 bind to multiple intronic sites, a plausible model is that their docking obstructs subsequent spliceosomal assembly. lncRNAs generally undergo less efficient splicing and consequently, they have higher levels of IR compared to protein coding genes (8,10). A study showed that lncRNA splicing strength correlates with 5′ ss strength and lncRNA Py-tract has higher thymidine content compared to protein coding genes (61). The reduced evolutionary pressure to maintain efficient splicing as they are not used as templates for protein synthesis, likely contributes to this pattern. This is also supported by their predominant nuclear localization (3,4).

Our data shows that *PURPL* transcripts can be either nuclear or cytoplasmic, and U2AF2 regulation of IR offers a mechanism of nuclear detention. Similar to this notion, the lncRNA *TUG1* remains in the nucleus due to intron retention (62), and the mouse lncRNA *pCharme* undergoes an IR event which renders it chromatin-bound through its association with MATR3 and PTBP1 (63). Another aspect of the effects of IR lies in the turnover of the transcripts where additional sequences added by the introns may cause lower transcript stability. In support of this, nuclear lncRNAs have been shown to be less stable than cytoplasmic ones (5), and *PURPL* also follows this rule. Notably, the human lncRNA *hFAST* is cytoplasmic and promotes pluripotency through WNT signaling, but the mouse homolog does not because of its nuclear localization (64).

Introns have been proposed to act as docking elements that tether lncRNAs to chromatin (4,63), and a recent study showed that nuclear speckles are associated with distinct classes of retained introns (65). This raises the question of what makes intron 2 important for speckle localization of *MALAT1*. One possibility is that this region recruits specific RBPs or adopts a structural configuration required for speckle association. Our data further indicates that intron 2 retention contributes to *MALAT1’s* role in cell migration, potentially because speckle localization is necessary to sustain this phenotype. Future studies are warranted to dig deeper into the underlying mechanism and to determine how intron retention shapes the function of *PURPL* and *MALAT1*.

Collectively, our findings reveal an expected function of U2AF2 in promoting intron retention that governs the subcellular localization and function of the lncRNAs *PURPL* and *MALAT1.* As intron retention continues to emerge as a regulated form of alternative splicing, the evidence presented here expands the framework for the mechanisms and biological consequences of intron retention.

## Materials and Methods

### Cell culture and treatment

HAP1, HCT116, HEK293T, HepG2, RPE1, SKHEP1, U2OS, WI38, and MDA-MB-231 cells were purchased from ATCC and maintained in Dulbecco’s Modified Eagle Medium (DMEM) (Gibco) supplemented with 10% (v/v) fetal bovine serum (Gibco) and 1% (v/v) PenStrep (Gibco) in a 5% CO_2_ atmosphere at 37°C. All cell lines were routinely confirmed to be free of mycoplasma using Venor GeM Mycoplasma Detection Kit (Millipore Sigma-Aldrich).

### CRASP-Seq *PURPL* screen

The *PURPL* minigene reporter, composed of exons 2 and 3 with the full native intronic sequence, was cloned into the CRASP-Seq genome-wide library as described previously (43). Reporter fragments were synthesized by TWIST Biosciences and assembled into the Eco32I-digested CRASP-Seq knockout library using NEBuilder (46). Library coverage (>250-fold) was maintained through parallel cloning reactions and electroporation into Endura competent cells followed by plasmid amplification and extraction.

Lentivirus was produced following established protocols (46). The pooled library was used to infect HAP1 and RPE1 cells expressing *Sp*Cas9 and opCas12a nucleases (46). at a multiplicity of infection (MOI) of ∼0.2. After puromycin selection (2 µg/mL for 48 hours), cells were pooled, induced with doxycycline to activate reporter expression and harvested 24 hours later for RNA analysis.

Total RNA was extracted, poly(A)+ RNA isolated, and cDNA synthesized using a U6-annealing primer. Spliced and intron-retaining reporter transcripts were amplified using exon- or intron-specific primers incorporating Illumina barcodes. Libraries were sequenced on the NovaSeq 6000 platform with a 25% PhiX spike-in and a 300-cycle kit. The sequencing strategy consisted of the following configuration: Read 1 (210 cycles), Index Read 1 (8 cycles), Index Read 2 (8 cycles) and Read 2 (85 cycles). Each sample yielded over 500 million total reads.

### CRASP-seq data analysis

CRASP-Seq data were processed using a custom pipeline (43). Guides, cell barcodes, and UMIs were retrieved using STAR, and alignment were managed with SAMTools. Reads containing complete hybrid guide, barcodes, UMI, and splicing outcomes were deduplicated prior to analysis. Splicing outcomes were quantified as percent intron retention (PIR): PIR = [IE / (IE + EE)] × 100, where IE represents intron retention reads, and EE represents exon-exon junction reads. Differential PIR (ΔPIR) was calculated as: ΔPIR = PIR_guide_ − PIR_intergenic_

The ΔPIR for each gene and for each guide separately is shown in Table S1. Guides with fewer than 10 reads were excluded. A ΔPIR threshold corresponding to 10% FDR (based on intergenic controls) defined significant hits. At the gene level, hits required FPKM ≥ 0.1, at least two qualifying guides, and ≥50% of guides exceeding the ΔPIR threshold in both technical replicates. Genes identified through combinatorial targeting that met these criteria were included as hits (Table S1). The heatmap and Venn diagram in Figure 1C and 1E display single-gene hits as well as collapsed gene hits resulting from combinatorial targeting.

### eCLIP-seq data and functional enrichment analysis

eCLIP-seq data for U2AF2, U2AF1, PTBP1, PRPF8, and SRSF1 were obtained from the ENCODE database (encodeproject.org) with the following identifiers: ENCFF989JBA (CTRL), ENCFF566CFJ (CTRL), ENCFF536AFD (U2AF2), ENCFF913WRH (U2AF2), ENCFF159SPZ (U2AF2), ENCFF368XEI (U2AF2), ENCFF542NOA (U2AF1), ENCFF651VRQ (U2AF1), ENCFF363UDO (PTBP1), ENCFF044RKN (PRPF8), and ENCFF327NVE (SRSF1). For visualization of RNA-seq data, the Integrated Genome Viewer (IGV) version 17 was used. The functional enrichment analysis was performed using g:Profiler (66) and the results are shown in the ‘gProfiler_Results’ tab in Table S1.

### siRNA transfections

For siRNA transfections, we used AllStars Negative Control siRNA (QIAGEN). U2AF2 siRNAs were purchased from Dharmacon (J-012380-18 and J-012380-19). U2AF1 siRNA was also purchased from Dharmacon (J-012325-10). Cells were reverse transfected with siRNA at a final concentration of 20 nM using Lipofectamine RNAiMAX (Invitrogen) in Opti-MEM I Reduced Serum Medium (Gibco) according to the manufacturer’s protocol. SKHEP1 cells were transfected for 72 hr before RNA harvesting. U2OS, HepG2 and WI38 cells were transfected for two rounds of 48 h + 72 h or 72 h + 72 h. For RNA FISH, SKHEP1, HCT116, and MDA-MB-231 cells were transfected for 72 hr. SKHEP1 expressing the *PURPL* Py-tract mutants were transfected for two rounds of 48 h and 24 h before harvesting, the cells were treated with 1 μg/mL doxycycline to induce the transgene expression. For RNA-seq, the cells were transfected for 72 h before harvested. For RNA stability assay, the cells were reseeded in 12-well plates 48 h after siRNA transfection and 24 h later they were treated with Act D for the indicated time points at 5 µg/mL before RNA extraction and subsequent assays. For RNA-IP, the cells were transfected for 72 h before harvested in 10 cm-plates.

### RNA extraction, RT-qPCR, semi-quantitative RT-PCR, and splicing gels

Total RNA was extracted using TRIzol (Invitrogen) according to the manufacturer’s protocol. cDNA was then made using iScript™ Reverse Transcription Supermix (Bio-Rad). For real-time qPCR, all reactions were carried out on StepOnePlus real-time PCR System (Applied Biosystems) using FastStart SYBR Green Master Mix (Millipore Sigma). *GAPDH* mRNA was used to normalize expression except for RNA-IPs in Figure 2B where *18S* was used for normalization, and the relative expression/enrichment of RNAs was calculated using the 2^-ΔΔCt^ method. RT-qPCR primer sequences for each gene are indicated in Table S3. For splicing gels, cDNA was used to amplify the region flanking the retained intron with primers in the flanking exons with the addition of one primer inside the intron as indicated in Figure S1B. In the case of *MALAT1*, primers in the flanking exons were utilized due to the short length of introns. For the assay of the effect of U2AF2 on the Py-tract mutants of *PURPL*, primers on the transgene were used instead of the endogenous *PURPL* exon 2 and 3. The sequences of the primers can be found in Table S5.

### RNA-IPs

For RNA-IP, two 10 cm-plates of SKHEP1 cells were lysed with RIPA buffer (Thermo Scientific) for 10 min on ice before spin down for 10 min at 4°C. The lysate was incubated with 1 μg of anti-U2AF2 (CST) or rabbit IgG (CST) in 500 μL total volume overnight at 4°C. The next day, 50 μL of Protein A/G Dynabeads (Pierce) were added to the mix and incubated for 2 h at 4°C. Beads were then washed 4 times with RIPA buffer and RNA was isolated using TRIzol. Equal volumes of extracted RNA were used for cDNA synthesis and RT-qPCR as described above.

### Immunoblotting

For immunoblotting, total cell lysate was prepared using RIPA buffer (Thermo Scientific) containing protease inhibitor cocktail (Roche). Protein concentration was determined using the BCA protein quantitation kit (Thermo Scientific). Cell lysate was loaded onto SDS-PAGE gel and transferred to a PVDF membrane (Thermo Scientific) using a semi-dry apparatus (Bio-Rad). The membrane was blocked in 5% skim milk (Millipore) in TBS containing 0.05% Tween 20. The following antibodies were used: anti-U2AF2 (1:1000, rabbit) (CST, 70471S), anti-GAPDH (1:10,000, rabbit) (CST, 2118L) and secondary anti-rabbit (1:5000) (CST, 7074S).

### Nuclear cytoplasmic fractionation

For nuclear cytoplasmic fractionation, SKHEP1 cells were harvested after trypsinization at 80-90% confluence. Cells were pelleted and resuspended in 500 μL RSB buffer (10 mM Tris-HCl, 100 mM NaCl, and 2.5 mM MgCl2) containing 40 μg/mL digitonin (Invitrogen), and 100 U of RNase Out. After incubation on ice for 10 min, cell viability was tested with trypan blue to make sure that the cells were permeabilized. The cytoplasmic fraction was harvested as supernatant after centrifugation at 1,500 × g for 5 min at 4°C. The nuclear pellet was resuspended in 100 µL RSB buffer. 100 µL of the cytoplasmic fractionation were transferred to a fresh tube and 900 µL of TRIzol was added as well as to the nuclear extracts for RNA extraction and subsequent cDNA synthesis and RT-qPCR.

### Cell proliferation assay

The cell proliferation assay was conducted as previously described (67). To measure proliferation, parental SKHEP1 cells and *PURPL*-CRISPRi cells overexpressing *PURPL* containing intron 2 were seeded at 1×10^^5^ cells per well in 6-well plates in the presence of 1 μg/mL doxycycline. After 3 days, the cell numbers and viability were measured. The cells were then passaged at 1×10^5^ and measured again at 6 days, and cumulative cell numbers were plotted.

### RNA-seq

The TruSeq Stranded mRNA libraries were pooled and sequenced on one NovaSeq6000 S1 flowcell using 2×101 cycles for pair-end run. The Real Time Analysis software (RTA v3.4.4) was used for processing raw data files, the Illumina bcl2fastq v2.20 was used to demultiplex and convert binary base calls and quality scores to fastq format. The sequencing reads were trimmed of adapters and low-quality bases using Cutadapt (v1.18). The trimmed reads were mapped to human reference genome (hg38) and Gencode annotation GENCODE_v30 GTF using STAR aligner (version 2.7.0f) with two-pass alignment option. RSEM (v1.3.1) was used for gene and transcript quantification based on GENCODE annotation file. Palantir Foundry was used for analysis of Differential Expression of Genes inside the secure NIH Integrated Data Analysis Platform (NIDAP). The DEG Analysis results are shown in Table S3.

### Iso-Seq

The libraries were prepared using PacBio standard Iso-seq library prep using SMRTbell prep kit 3.0. The libraries were sequenced on PacBio Sequel II platform using PacBio v2.0 chemistry.

PacBio SMRTlink raw subreads were converted into HiFi circular consensus sequences (CCS) with minimum 3 passes. PacBio IsoSeq v3 pipeline was used to demultiplex the barcodes and remove primers. Additional refine steps included trimming poly-A tails and removing concatemers to generate Full Length Non-Concatemer (FLNC) reads. The FLNC reads were used to map to human reference (hg38) using Minimap2 software to generate alignment bam files. Hierarchical clustering and merging steps were performed to obtain consensus isoforms and the full-length (FL) consensus sequences. The high-quality FL transcripts were mapped to the reference genome by using the minimap2 software. Isoform classification and quality control were done using SQANTI3 software and Illumina short-read gene expression data was combined with PacBio Iso-seq for transcript quantification. The Iso-Seq Analysis results are shown in Table S4. The resulting bam files from the Iso-seq analysis were used for plotting and visualization with the R package Gviz (https://bioconductor.org/packages/release/bioc/html/Gviz.html).

### IRFinder

IRFinder (52) was used to identify intron retention events that changed significantly on U2AF2 knockdown. A docker image was downloaded from Docker Hub (docker://cloxd/irfinder:2.0.1). The reference was built using pre-downloaded resources (IRFinder BuildRefProcess), using Ensembl release 92 and the 100 bp mappability file provided by IRFinder. The Build-BED-refs.sh script was adjusted so that transcript biotype was not restricted to “processed_transcript” or “protein_coding”. Intron retention was quantified using IRFinder in FastQ mode with adapter trimming. Differential intron retention events between siU2AF2 vs. siCTRL were identified with IRFinder Diff using DESeq2 with the default IR ratio ≥ 0.05 in at least one sample and no warning filters, which profiled 16,859 unique introns.

### Construct generation

For Py-tract mutants, gene fragments were purchased from Twist Biosciences and cloned into the pLCHKO-puro vector containing the WT sequences of *PURPL* (Figure 1A), between sites for NseI and AfeI. The *PURPL* transcript containing intron 2 was cloned into the pCW57 vector (Hygro) (Addgene #80922) in SalI and MluI sites. *MALAT1* transcripts were cloned into the pCW57 vector (BSD) (Addgene #80921) in SalI and MluI sites. The *PURPL* insert was generated by Twist Biosciences and the *MALAT1* sequence was amplified using Phusion DNA polymerase (NEB). For deletion mutants, we used the Q5® Site-Directed Mutagenesis Kit (NEB). The primers used for cloning are listed in Table S5.

### Generation of *MALAT1* KO cell lines

For HCT116 *MALAT1* knockout, we used integration by non-homologous end joining, which was accomplished by introducing a simultaneous double-strand break in genomic DNA and in the targeting vectors (pCMV-Puro and pCMV-Hygro) (68,69). Plasmids encoding spCas9 and sgRNAs were obtained from Addgene (Plasmids #41815 and #47108). Oligonucleotides for construction of sgRNAs were obtained from Integrated DNA Technologies, hybridized, phosphorylated and cloned into the sgRNA plasmid or targeting vector using BbsI sites (70). Target sequences for sgRNAs are provided in Table S5. These vectors were transfected into HCT116 using Lipofectamine 2000 following recommendations from the manufacturer. Three days after transfection, the cells were selected with Puromycin (0.5 μg/mL) and Hygromycin (100 μg/mL) to generate clonal populations. Genomic DNA from each clone was isolated using DNEasy Blood and Tissue Kit (Qiagen). PCRs to detect integration of the targeting vector at the target site were performed using KAPA2G Robust PCR kits (Kapa Biosystems) according to the manufacturer’s instructions. A typical reaction contained 20-100 ng of genomic DNA in Buffer A (5 µL), Enhancer (5 µL), dNTPs (0.5 µL), primers forward (PINCR Det FP, 1.25 µL) and reverse (Targeting vector Det RP, 1.25 µL) and KAPA2G Robust DNA Polymerase (0.5 U). The DNA sequences of the primers for each target are provided in Table S5. PCR products were visualized in 2% agarose gels and images were captured using a ChemiDoc-It2 (UVP).

To generate MDA-MB-231 whole-locus and intron 2 *MALAT1* knockout clones, we used the CRIPSR/Cas9 system by transfecting the cells with gRNAs and purified Cas9. 2 µL of each 30 µM gRNA was mixed with 1 µL (∼5 μg) of purified Cas9 and 100 µL of 4D-Nucleofector® X SE buffer SE from the Cell Line Kit (Lonza) supplemented with 18.5 µL of SE supplement per 100 µL total buffer volume. The mixture was incubated at room temperature for 10 min and 1×10^6 cells were resuspended in nucleofection cuvettes. Nucleofections were carried out using the 4D-Nucleofector® System (Lonza). The cells were transferred to a 6-well plate, and they were single-cell sorted 48 h later. The clones were grown for 3-4 weeks before genomic DNA extraction and genotyping with PCR using the primers in Table S5.

### *PURPL* and *MALAT1* overexpression and transduction

*PURPL* intron 2 constructs were transduced in previously generated *PURPL*-CRISPRi SKHEP1 cells (11). *MALAT1*-WT and deletion mutants were transduced in the two HCT116-*MALAT1* KO clones. For lentivirus generation, HEK293T cells were seeded at 2×10^5^ cells/well onto 6-well plates, and 1,200 ng of each construct was transfected with lentiviral package vectors using Lipofectamine 2000 (Life Technologies, Invitrogen) according to the manufacturer’s protocol. Virus-containing media were harvested at 48 and 72 h post-transfection and added to cells for 24 h. Transduced cells were selected with 500 μg/mL hygromycin for 2 weeks (SKHEP1) or 10 μg/mL Blasticidin (HCT116).

### RNA-FISH, IF and Quantification of FISH

The *PURPL* intron probe set was custom-designed using the Stellaris Probe Designer, tagged with Quasar 570 dye and purchased from Stellaris. The *MALAT1* smFISH probe set with Quasar 670 dye was purchased from Stellaris (Cat# VSMF-2211-5). For *PURPL* intron smRNA-FISH, SKHEP1 cells were seeded on coverslips and cultured for 24 hours at 37°C with 5% CO_2_ before inducing DNA damage by incubating the cells with 2 mM hydroxyurea for 24 hours (or regular growth media for control). For *MALAT1* smRNA-FISH, HCT116 and MDA-MB-231 *MALAT1* KO cells were seeded on coverslips and cultured for 24 hours at 37°C with 5% CO_2_. *MALAT1* WT, *MALAT1* del-1, *MALAT1* del-2, *MALAT1* del1+2, or empty vector expression was induced by incubating the cells with 1 µg/mL doxycycline for 48 hours. The cells that were transfected with siRNA for U2AF2 were seeded on coverslips at the time of transfection. Cells were fixed with 4% PFA for 15 min at room temperature, permeabilized with 0.5% Triton X-100, and washed with washing buffer (10% formamide, 2XSSC) for 5 min. Probe was added to hybridization buffer (10% formamide in Stellaris hybridization buffer Cat# SMF-HB1-10) at a final concentration of 125 nM. Hybridization was done in a humidified chamber in the dark overnight at 37°C. After hybridization, the coverslips were washed twice with wash buffer for 30 min at 37°C and then post-fixed with 4% PFA for 10 min at room temperature. The coverslips were blocked with 5% normal goat serum and then incubated with SON antibody (Sigma Cat# HPA023535). Primary and secondary antibodies were diluted in 1% normal goat serum. Antibody incubations were done at room temperature for 1 h and washed with PBS. DNA was counterstained by DAPI. The coverslips were then washed with 4XSSC for 5 min at room temperature and mounted in VectaShield Antifade Mounting Medium (Vector Laboratories). Images were taken using DeltaVision microscope (GE Healthcare) equipped with 60 X/1.42 NA oil immersion objective (Olympus) and CoolSNAP-HQ2 camera. To quantify the speckle to nucleoplasm signal intensity ratio of *MALAT1* in Figures 5B, 7B, 7D, and S8B ImageJ was used to create nucleus masks based on DAPI and speckles mask based on SON. The signal intensity of *MALAT1* in the speckles mask was divided by the *MALAT1* signal intensity in the nucleus-minus-speckles area. For the quantification of Figures 6B, 7E and S10B, the cells with *MALAT1* FISH signal showing prominent speckle patterns similar to SON were classified as “Speckle” pattern, and the cells with a diffused *MALAT1* FISH signal were classified as “Diffused”. Only *MALAT1*-positive cells were counted for quantification.

### Transwell cell migration assays

For the transwell assay, WT and KO MDA-MB-231 clones were seeded at 10×10^5^ cells per well in 12-well plates. 24 h later, the media were removed and replaced with media containing 1% FBS. After 5 h, the cells were trypsinized and seeded in the top chamber of the transwell plates (Corning, Catalog#354578) at 25,000 cells per well at a volume of 100 μL. The lower chamber of the transwell is filled with 700 μL of media containing 10% FBS. The cells were incubated for 24 h and the transwell were removed from the plates. The non-migrated cells were scraped with a cotton swab, and the migrated cells were fixed with ice-cold methanol for 5 min and stained with 0.05% crystal violet blue. The migrated cells were counted under a light microscope.

### Statistics

Statistics were performed using the Student’s t-test.

## Data availability

The Iso-Seq data files have been deposited at GEO with accession number GSE288684 and reviewer token: ktcnasayrbgnjev (https://www.ncbi.nlm.nih.gov/geo/query/acc.cgi). The RNA-Seq data files have been deposited at GEO with accession number GSE288685 and reviewer token: qvwhuawixxqjnml (https://www.ncbi.nlm.nih.gov/geo/query/acc.cgi).

## Acknowledgments

This research was supported by the Intramural Research Program (A.L.) of the National Cancer Institute (NCI), Center for Cancer Research (CCR) (project numbers ZIA BC011646, ZIA BC012019), NIH. KVP laboratory is supported by grants from NIH (R01-GM132458), Cancer center at Illinois seed grant, ARPA-H, and National Science Foundation (NSF) center for quantitative cell biology. YJS is supported by NIH T32EB019944. We thank the CCR Genomics Core, CCR, NCI, Bethesda, MD for its valuable assistance with Sanger and Nanopore sequencing. We also thank the Sequencing Facility, CCR, NCI, Frederick, MD for performing the RNA sequencing assays. The contributions of the NIH author(s) were made as part of their official duties as NIH federal employees, are in compliance with agency policy requirements, and are considered Works of the United States Government. However, the findings and conclusions presented in this paper are those of the author(s) and do not necessarily reflect the views of the NIH or the U.S. Department of Health and Human Services.

## Author Contributions

IG, CN, YJS, CCH, SK, RP, XLL, RK, RS, and PPZ conducted experiments. IG, AKB, EP, YZ, BS, TB, XW, TGP, KP, and AL did data analysis. IG and AL wrote the manuscript. NC, KP, and TGP edited the manuscript.

