## Supplemental Figures for "Intron Retention Controls Localization of lncRNAs *PURPL* and *MALAT1* to Promote Cell Proliferation and Migration"

### Figure S1

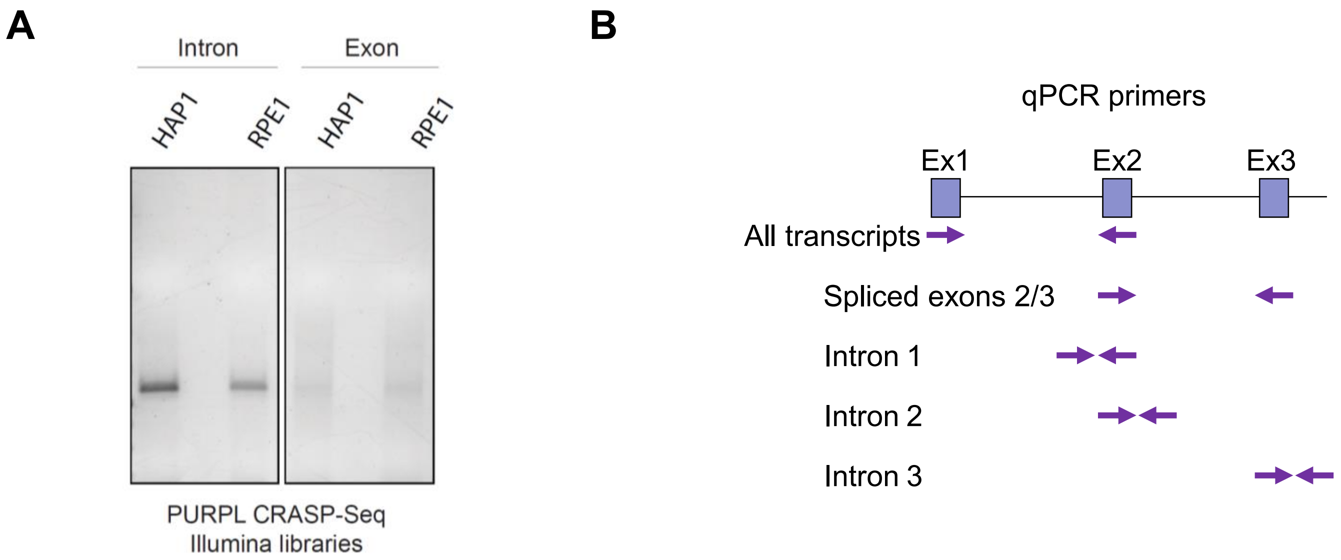
A
Intron
Exon
HAP1
RPE1
HAP1
RPE1
PURPL CRASP-Seq
Illumina libraries
B
qPCR primers
Ex1
Ex2
Ex3
All transcripts
Spliced exons 2/3
Intron 1
Intron 2
Intron 3

**Figure S1. Screen validation. (A)** Gel showing the intensity of the Illumina libraries before sequencing indicating that most of the transcripts contain the intron. **(B)** Schematic showing where the primer pairs are designed for RT-qPCR to distinguish intron 2-containing and spliced transcripts of *PURPL*. The primer pair designed on exon 1 recognizes all transcripts whereas the rest are specific for transcripts where exons 2 and 3 are spliced together or for transcripts containing introns.

#### Figure S2

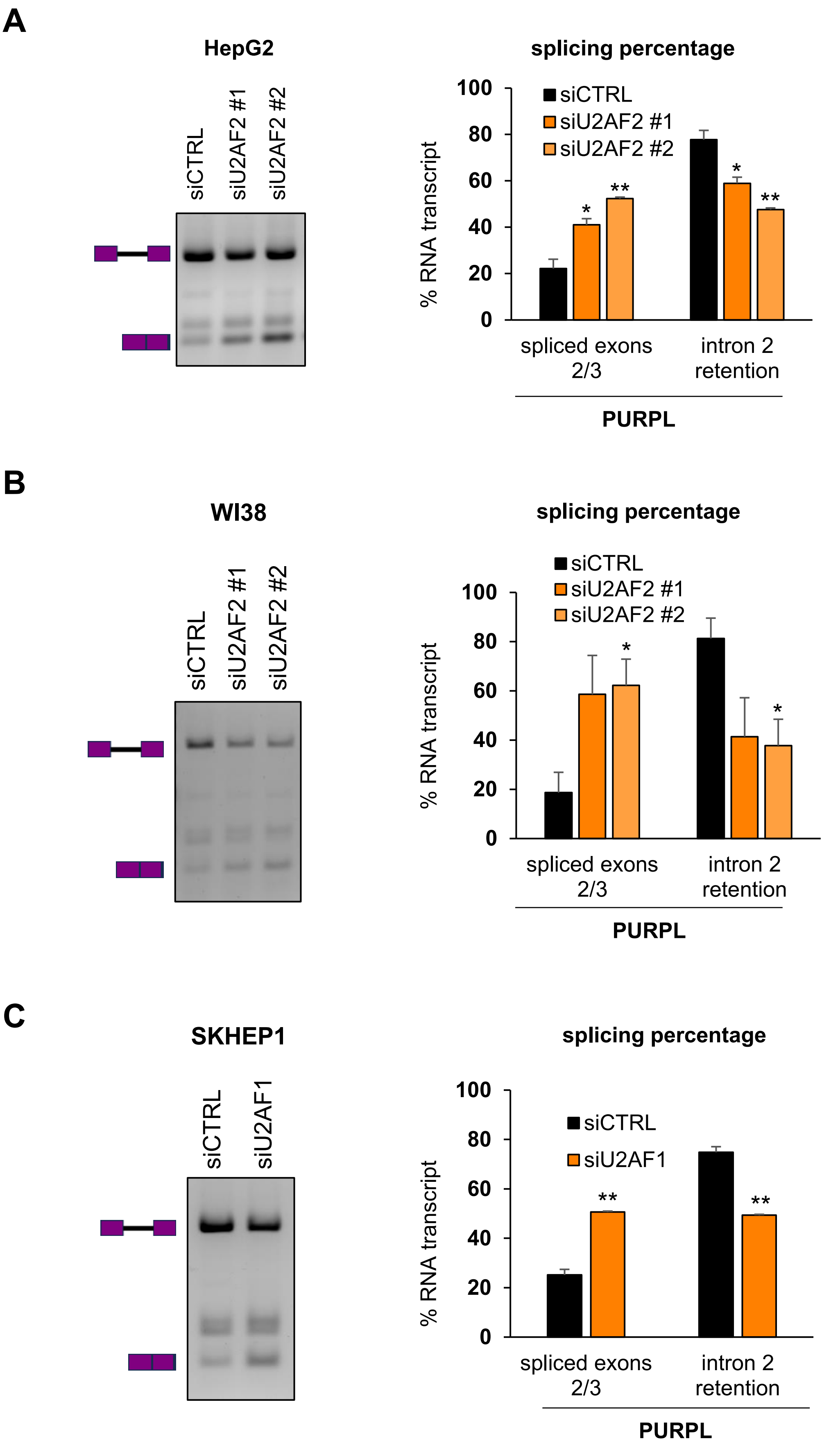

A
HepG2
siCTRL
siU2AF2 #1
siU2AF2 #2
splicing percentage
% RNA transcript
siCTRL
siU2AF2 #1
siU2AF2 #2
spliced exons 2/3
intron 2 retention
PURPL
B
WI38
siCTRL
siU2AF2 #1
siU2AF2 #2
splicing percentage
% RNA transcript
siCTRL
siU2AF2 #1
siU2AF2 #2
spliced exons 2/3
intron 2 retention
PURPL
C
SKHEP1
siCTRL
siU2AF1
splicing percentage
% RNA transcript
siCTRL
siU2AF1
spliced exons 2/3
intron 2 retention
PURPL

**Figure S2. Detection of *PURPL* transcripts in RT-PCR in multiple cell lines. (A)** and **(B)** RT-PCR for *PURPL* transcripts upon knockdown of U2AF2 with 2 different siRNAs in **(A)** HepG2, and **(B)** WI38 cells. **(C)** RT-PCR for *PURPL* transcripts upon knockdown of U2AF1 in SKHEP1 cells. The schematics next to the gel indicate the expected products of the intron-retained and spliced isoforms, respectively. Quantitation of the gel bands is shown in the graphs on the right. Error bars represent the SD of 2 experiments. *p<0.05, **p<0.01.

### Figure S3

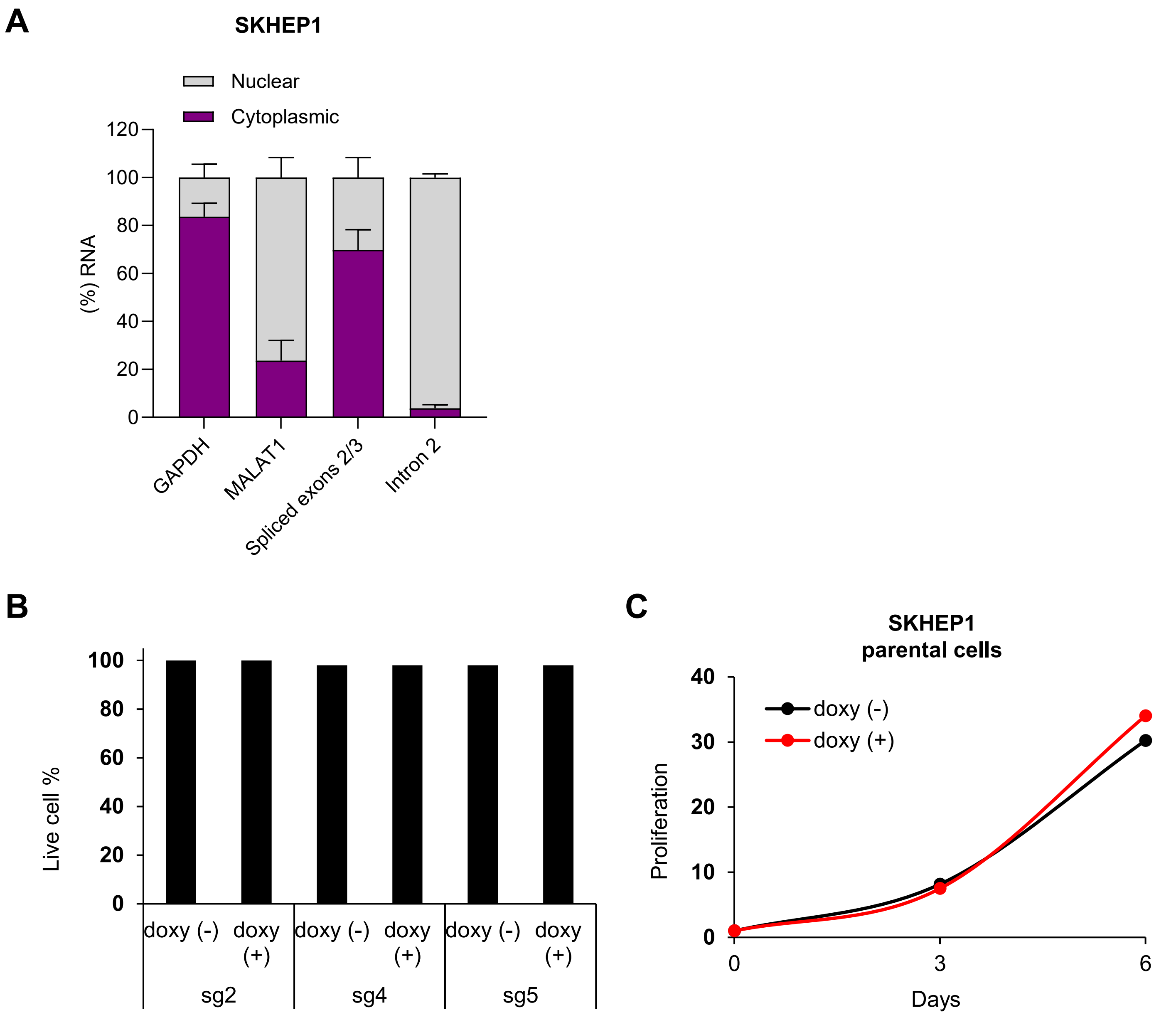

A
SKHEP1
Nuclear
Cytoplasmic
(%) RNA
120
100
80
60
40
20
0
GAPDH
MALAT1
Spliced exons 2/3
Intron 2
B
Live cell %
100
80
60
40
20
0
doxy (-)
doxy (+)
sg2
sg4
sg5
C
SKHEP1
parental cells
doxy (-)
doxy (+)
Proliferation
40
30
20
10
0
3
6
Days

**Figure S3. Intron 2-retained *PURPL* is localized to the nucleus. (A)** RT-qPCR shows the distribution of *PURPL* transcripts after nuclear cytoplasmic fractionation from SKHEP1 cells. The location of the primers in transcripts is shown in Figure S1C. When intron 2 is retained, *PURPL* is exclusively nuclear while the isoform with the spliced intron out is mostly cytoplasmic. *GAPDH* was used as a cytoplasmic and *MALAT1* as a nuclear control. The error bars show standard deviations of 3 independent experiments. **(B)** Proliferation assay of parental SKHEP1 cells to validate that doxycycline treatment does not affect cell proliferation. The cells were treated with 1 μg/mL doxycycline and cell proliferation was monitored at 3 and 6 days. **(C)** Graph showing the live cell percentage at the end of the proliferation assay (Fig. 2E) indicating that the cell viability was not affected.

**Figure S4**

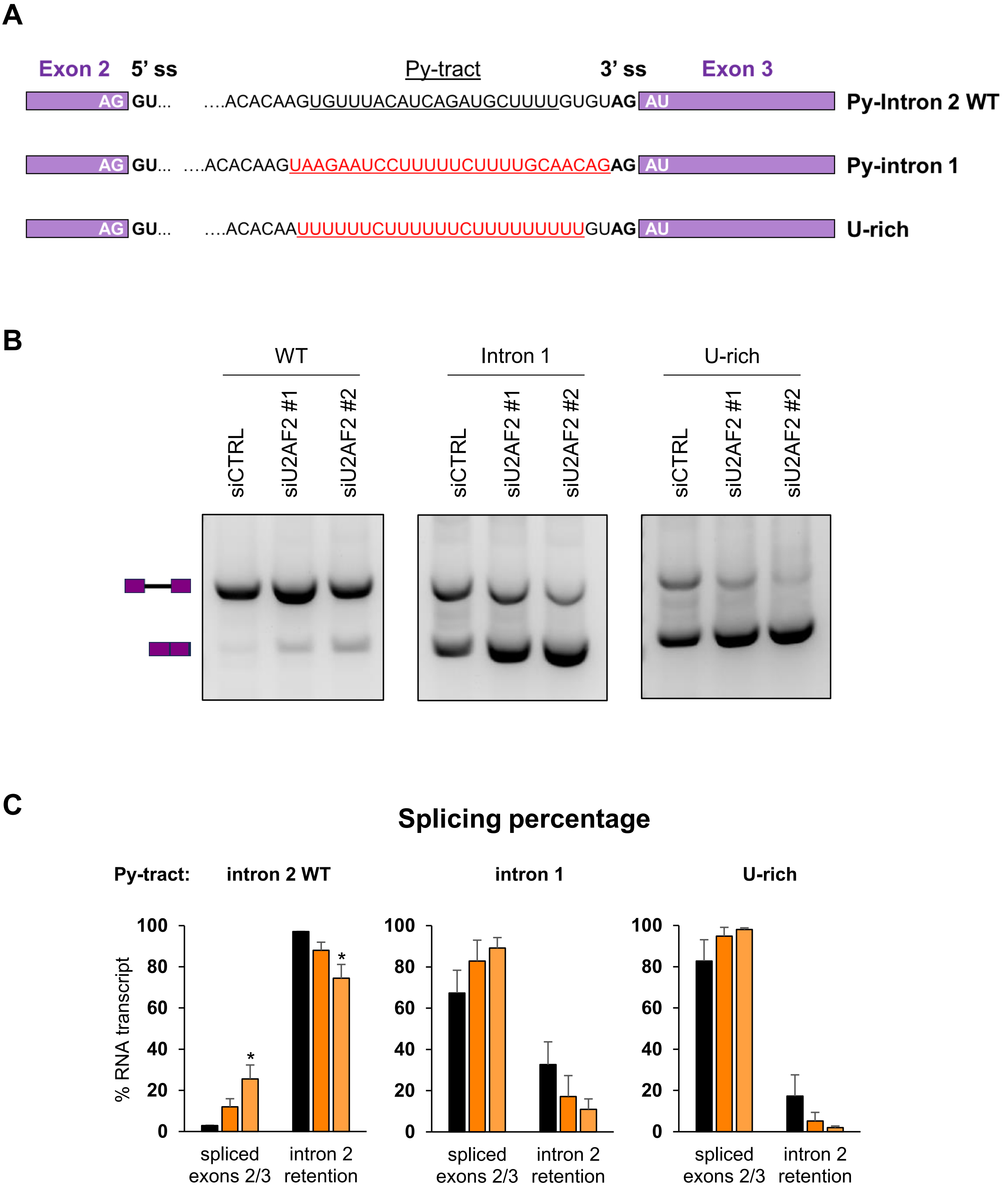

A
Exon 2
5’ ss
Py-tract
3’ ss
Exon 3
AG GU… ….ACACAAGUGUUUACAUCAGAUGCUUUUGUGUAG AU
Py-Intron 2 WT
AG GU… ….ACACAAGUAAGAAUCCUUUUUCUUUUGCAACAGAG AU
Py-intron 1
AG GU… ….ACACAAUUUUUUCUUUUUUCUUUUUUUUUGUAG AU
U-rich
B
WT
Intron 1
U-rich
siCTRL
siU2AF2 #1
siU2AF2 #2
C
Splicing percentage
Py-tract:
intron 2 WT
intron 1
U-rich
% RNA transcript
100
80
60
40
20
0
*
spliced exons 2/3
intron 2 retention

**Figure S4. Py-Tract is important for *PURPL* IR regulation by U2AF2. (A)** Panels showing the sequences of the Py-tract of the 3 pLCHKO constructs used as shown in Figure 1A to study the effect of the Py-tract in *PURPL* intron 2 retention. The original sequence is indicated in the top construct where the Py-tract is underlined (WT). The Py-tract was replaced by the one from intron 1 of (intron 1), or with a stretch of Us (U-rich). **(B)** RT-PCR upon knockdown of U2AF2 with 2 different siRNAs in SKHEP1 cells for constructs with WT or mutated Py-tract in intron 2. The primer triplet used specifically recognizes the transgene and intron 2 (Fig. 2C). The schematics next to the gel indicate the expected products of the intron-retained and spliced isoforms, respectively. **(C)** Bar graph with quantitation of the gel bands in **(B)**. Error bars represent standard deviations from 2 independent experiments. *$p<0.05$.

### Figure S5

A

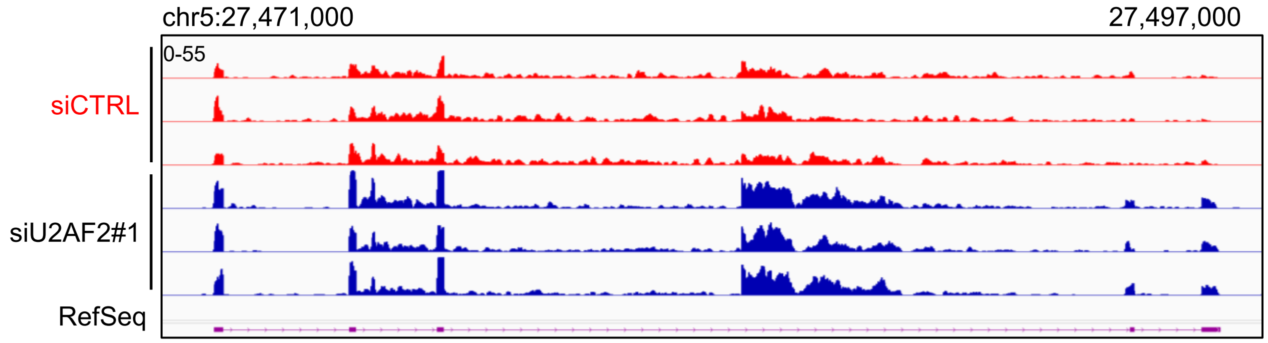


B

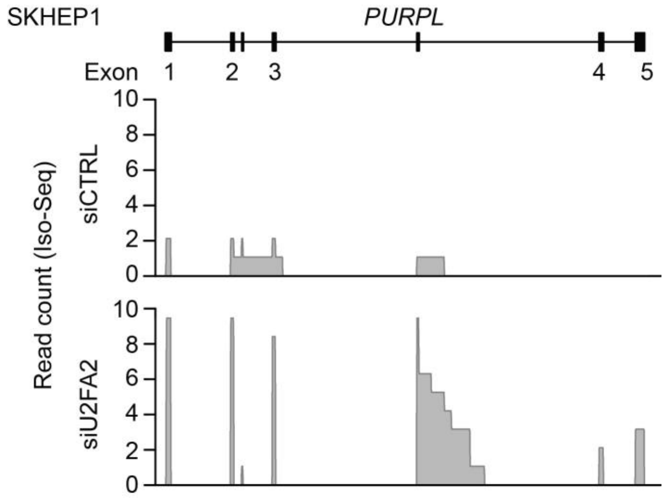


C

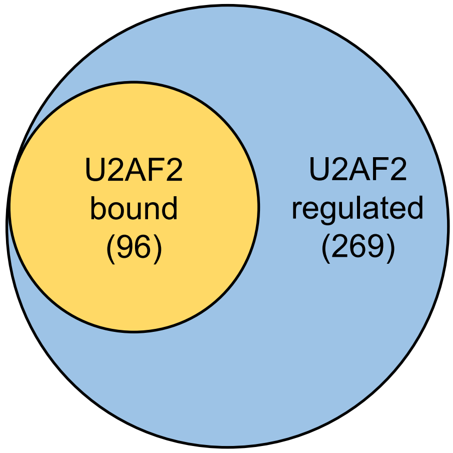

**Figure S5. U2AF2 binds to intron 2-containing *PURPL* transcripts and promotes intron 2 retention. (A)** IGV snapshot of RNA-Seq read coverage at the *PURPL* locus in SKHEP1 cells in 3 siCTRL (red) and 3 siU2AF2#1 (blue) samples. **(B)** Gviz coverage plots of long-read transcript variants from siCTRL (top) or siU2AF2#1 (bottom) SKHEP1 cells identified using Iso-seq. The Y axis represents coverage, indicating the total full-length counts for the indicated region within the transcript. The canonical transcript structure provided as a reference. **(C)** Venn diagram showing the number of intron retention events regulated by U2AF2 along with the number of these introns where U2AF2 binds to the intron (inside or in close proximity) as analyzed from eCLIP-seq data.

### Figure S6

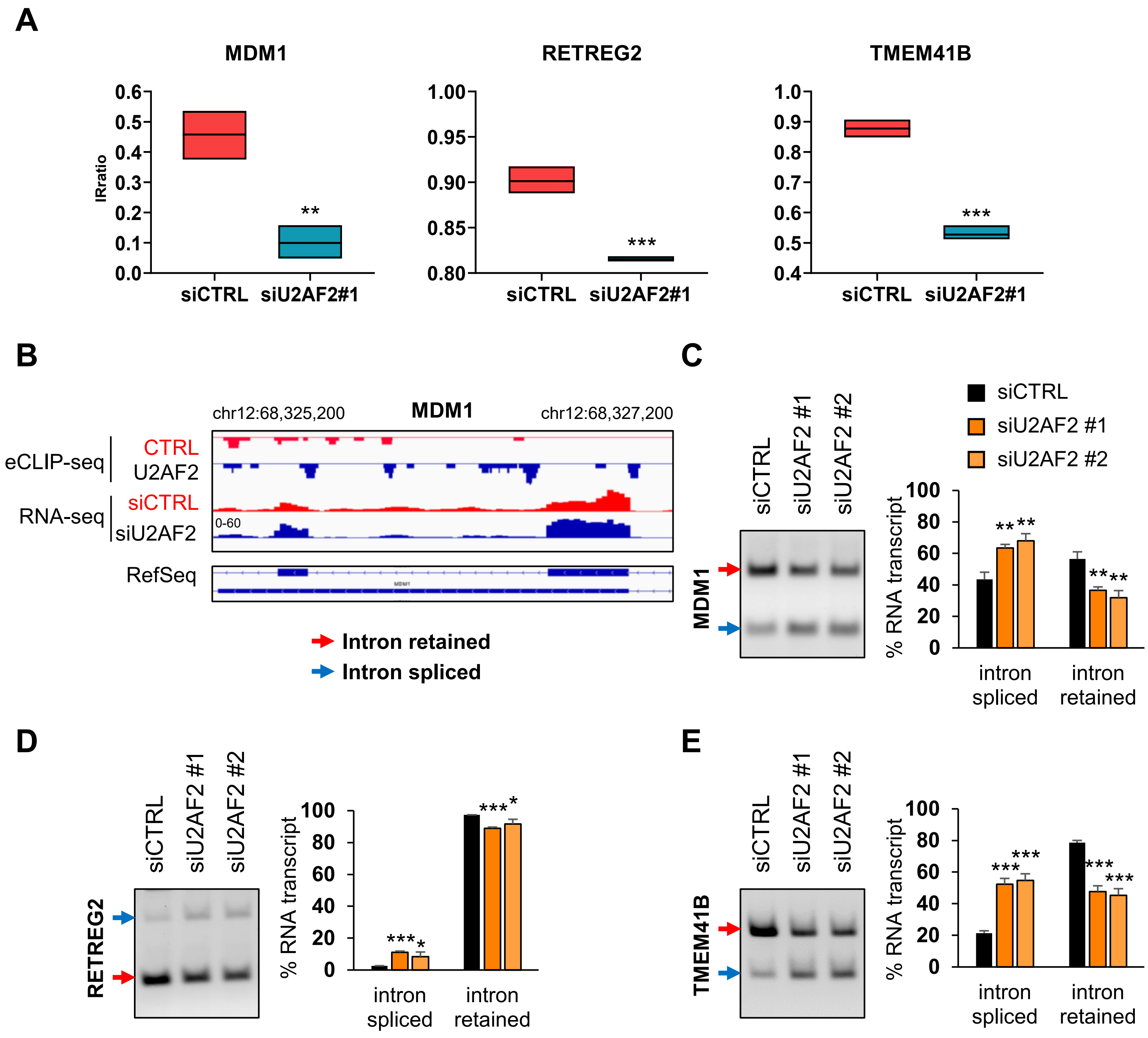


**Figure S6. U2AF2 directly regulates multiple IR events. (A)** Floating bar plot showing the IR ratio in genes *MDM1*, *RETREG2*, and *TMEM41B* in siCTRL and siU2AF2#1 samples as analyzed with the IRFinder algorithm. **(B)** IGV snapshot of Control (red) and U2AF2 eCLIP data (blue) showing binding sites and enrichment of U2AF2 on or around the retained intron of *MDM1*, and RNA-seq in siCTRL (red) and siU2AF2 (blue) samples in SKHEP1 cells (bottom). The annotated locus by RefSeq showing the retained intron is also indicated. eCLIP data were analyzed from encodeproject.org. The gene is expressed from the negative strand. **(C)**, **(D)**, and **(E)** *Left*: RT-PCR for the indicated transcripts using primer triplets upon knockdown of U2AF2 with 2 different siRNAs in SKHEP1 cells. The bands corresponding to intron-retained and intron-spliced variants are indicated with red and blue arrows, respectively. *Right*: Bar graph with quantitation of the gel bands. Error bars represent standard deviations from 3 independent experiments. *p<0.05, **p<0.01, ***p<0.001.

### Figure S7

**A**

Intron 2

…CACTTTTAGGCCATCAAACTCCAGTCTGCACCTGGAGCCTTGGATGATGGCCTTTTTACTGG
GAACCCTGAAATAGGCCTCTGAGGGAAATCTGATTGCCATTTTCCCAAAACAGCTTCCCTGACA
ATAGGAAGCAATTAAGATTGGTCTTCATCCTTATCTTTATCCTTATTCTAATGGCAGTTAGAGGT
ACTTCTTTAGAGGGGGGAATGAGACAGCCAAATGCCTAGGCAGATAAAAAGGTGTACCTGGAA
AATTTCCAACCCGTCACACAAGTGTTTACATCAGATGCTTTTGTGTAGATGAGGGAAACTT

**B**

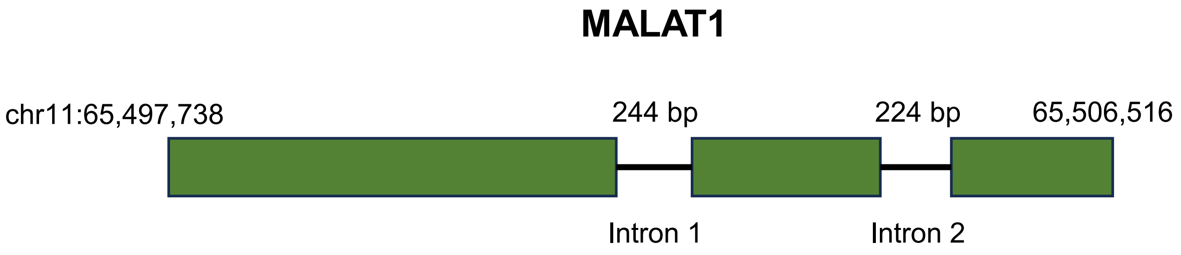


**C**

MALAT1 Intron 1

GTATTATCAAGTAAGATTCTATTTTCAGTTGTGTGTAAGCAAGTTTTTTTTTAGTGTAGGAGAAAT
ACTTTTCCATTGTTTAACTGCAAAACAAGATGTTAAGGTATGCTTCAAAAATTTTGTAAATTGTTT
ATTTTAAACTTATCTGTTTGTAAATTGTAACTGATTAAGAATTGTGATAGTTCAGCTTGAATGTCT
CTTAGAGGGTGGGCTTTTGTTGATGAGGGAGGGGAAACTTTTTTTTTTTCTATAGACTTTTTTCA

MALAT1 Intron 2

TGTATTTTTAGGTAAAATGCTTTTTGTTCATTTCTGGTGGTGGGAGGGGACTGAAGCCTTTAGTC
TTTTCCAGATGCAACCTTAAAATCAGTGACAAGAAACATTCCAAACAAGCAACAGTCTTCAAGAA
ATTAAACTGGCAAGTGGAAATGTTTAAACAGTTCAGTGATCTTTAGTGCATTGTTTATGTGTGGG
TTTCTCTCTCCCCTCCCTTGGTCTTAATTCTTACATGCAGGAACACTCAG

**Figure S7. *PURPL* and *MALAT1* intron retention events sequences. (A)** *PURPL* intron 2 sequences proximal to the 3′ss (top). **(B)** Diagram showing the gene structure of *MALAT1* with its coordinates. **(C)** *MALAT1* intron 1 (top) and intron 2 (bottom) sequences are indicated. In **(A)** and **(C)**, red font indicates exonic sequences while black indicates intronic. The underlined sequences are the potential Py-tracts.

### Figure S8

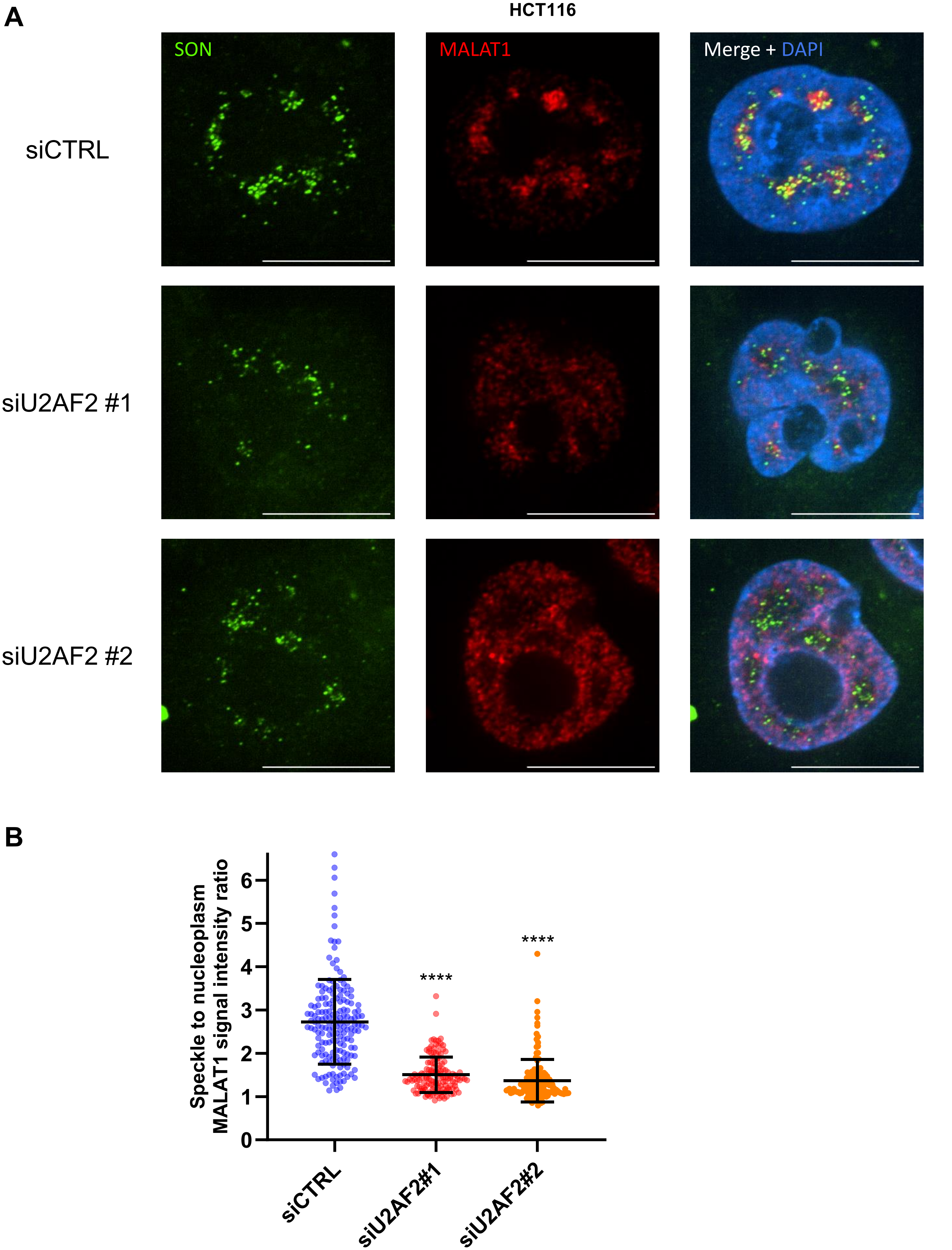
A
HCT116
SON
MALAT1
Merge + DAPI
siCTRL
siU2AF2 #1
siU2AF2 #2
B
Speckle to nucleoplasm MALAT1 signal intensity ratio
0
1
2
3
4
5
6
****
****
siCTRL
siU2AF2#1
siU2AF2#2

**Figure S8. U2AF2 promotes localization of *MALAT1* to nuclear speckles in HCT116. (A)** RNA-FISH images for *MALAT1* and Immunofluorescence images for SON is shown upon transfection of HCT116 cells with siCTRL or siU2AF2. *MALAT1* is enriched in nuclear speckles in the siCTRL but not upon U2AF2 knockdown. **(B)** Quantitation of the speckle to nuclear plasma *MALAT1* signal ratio in the three replicates in panel **(A)**. ****$p<0.0001$.

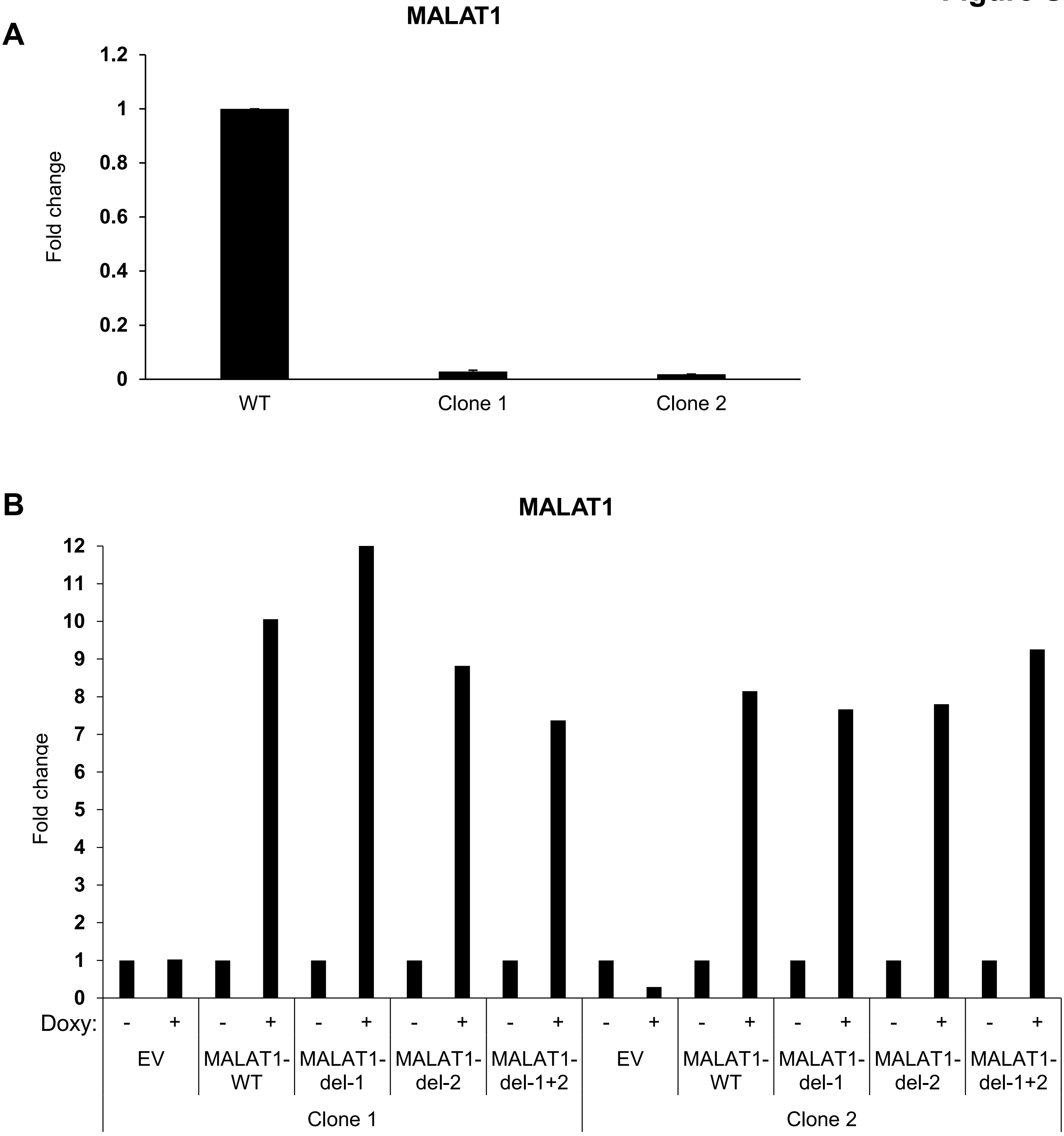


**Figure S9. Re-introduction of *MALAT1* transcripts in *MALAT1* KO cells. (A)** RT-qPCR showing *MALAT1* depletion in 2 different HCT116 clones compared to a WT clone. **(B)** RT-qPCR for *MALAT1* transcripts after 48 hr of 1 µg/mL doxycycline treatment in comparison to no treatment for each *MALAT1* transcript in both knockout clones. EV: Empty vector control.

### Figure S10

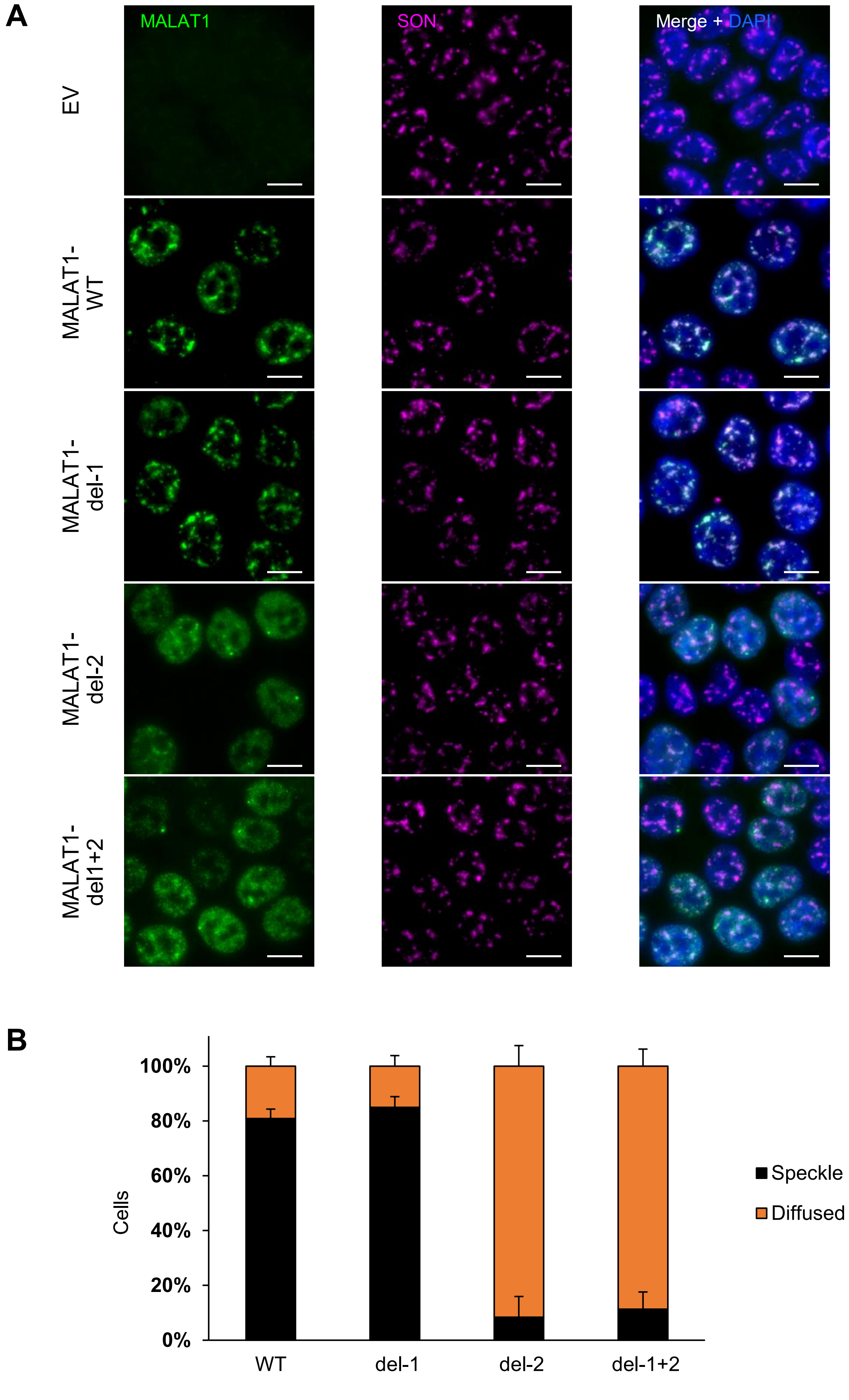
A
MALAT1
SON
Merge + DAPI
EV
MALAT1-WT
MALAT1-del-1
MALAT1-del-2
MALAT1-del1+2
B
Cells
100%
80%
60%
40%
20%
0%
WT
del-1
del-2
del-1+2
Speckle
Diffused

**Figure S10. Intron 2 of *MALAT1* dictates its speckle localization in more than one *MALAT1* KO clone. (A)** RNA FISH images for *MALAT1* and Immunofluorescence images for SON in a HCT116 *MALAT1*-KO clone (clone 2) where Empty Vector (EV) or constructs that overexpress *MALAT1* transcripts were re-introduced. Transcripts had full-length (WT), intron 1 deleted (del-1), intron 2 deleted (del-2) or both introns deleted simultaneously (del-1+2). The cells were first treated with 1 μg/mL doxycycline for 48 hr to induce *MALAT1* expression. Scale bar is 10μm. **(B)** Graphs showing quantification of the percentage of cells where *MALAT1* transcripts are enriched in speckles or being diffused in HCT116 cells clone 2 related to **(A)**. Error bars represent standard deviations from 2 independent experiments.

### Figure S11

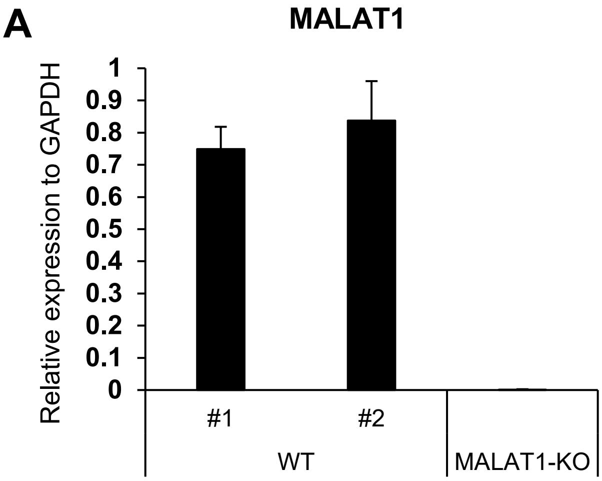
A
MALAT1
Relative expression to GAPDH
1
0.9
0.8
0.7
0.6
0.5
0.4
0.3
0.2
0.1
0
#1
#2
WT
MALAT1-KO

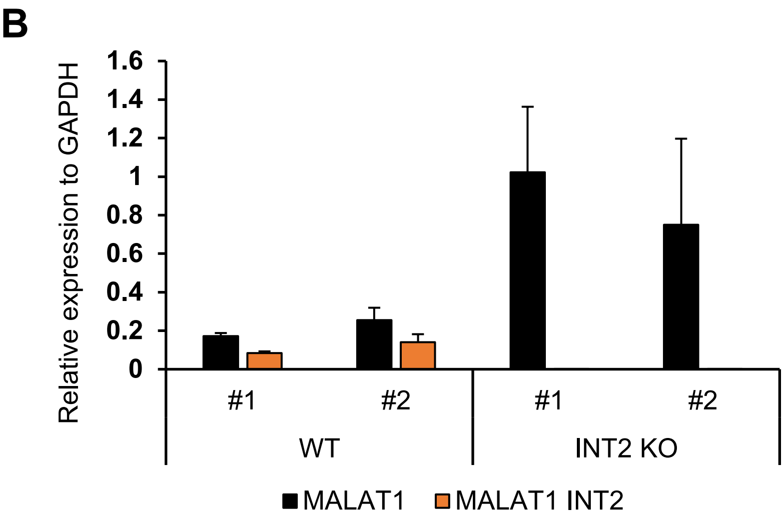
B
Relative expression to GAPDH
1.6
1.4
1.2
1
0.8
0.6
0.4
0.2
0
#1
#2
#1
#2
WT
INT2 KO
MALAT1
MALAT1 INT2

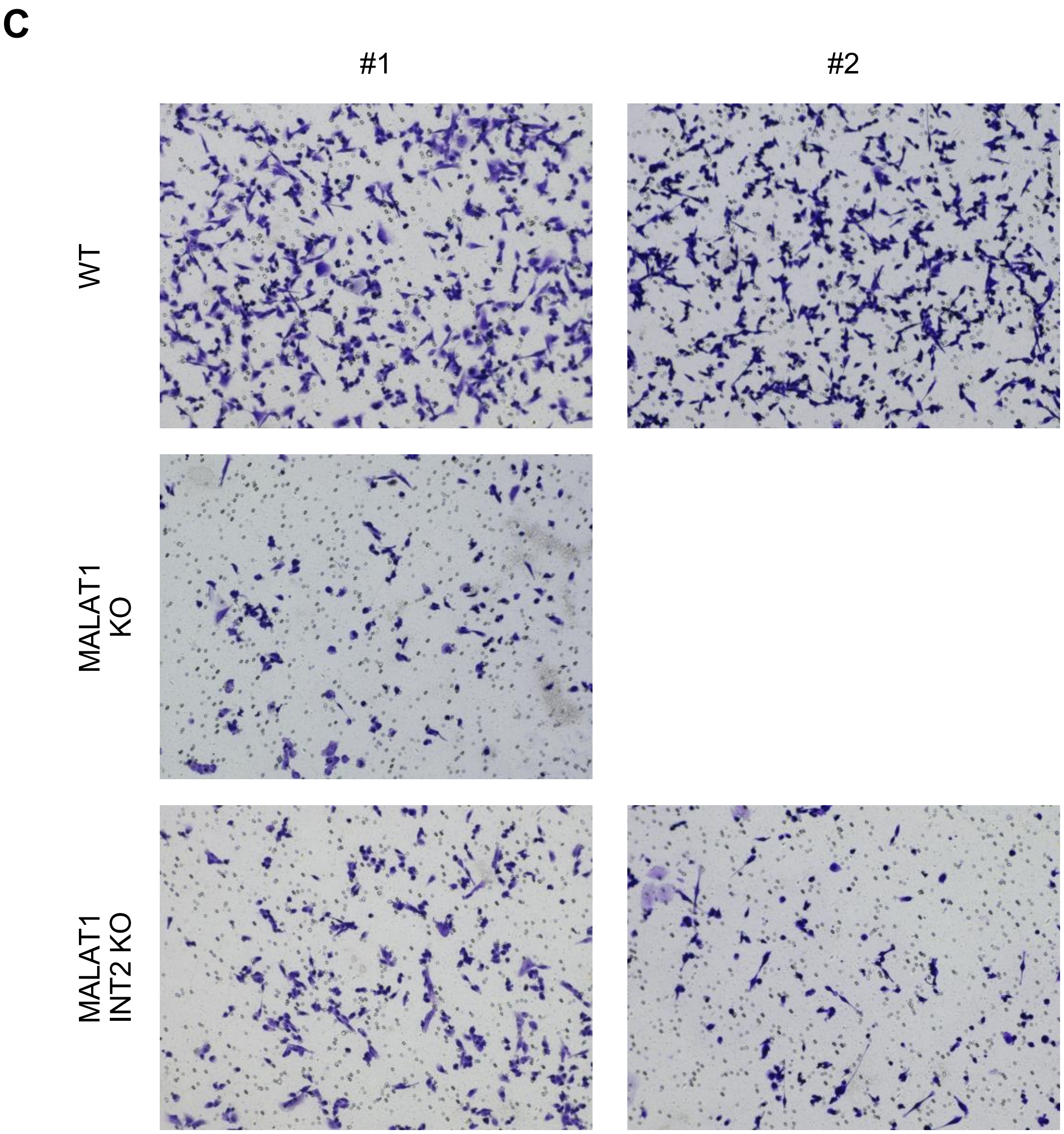
C
#1
#2
WT
MALAT1 KO
MALAT1 INT2 KO

**Figure S11. Deletion of intron 2 of *MALAT1* leads to decreased migration potential of breast cancer cells.** **(A)** RT-qPCR showing depletion of *MALAT1* in an MDA-MB-231 *MALAT1-KO* clone compared to 2 WT clones. **(B)** RT-qPCR showing the expression of *MALAT1* transcripts upon intron 2 deletion in 2 WT and 2 deletion clones. 2 pairs of primers were used, one that recognizes all *MALAT1* transcripts and one only the transcripts that include intron 2 (Table S5). The graphs in **(B)** and **(C)** show the levels of expression compared to *GAPDH,* N=2. **(C)** Representative pictures showing MDA-MB-231 cells that have migrated after 24 hr in a transwell assay. Each picture corresponds to one clone (WT, *MALAT1*-KO, or *MALAT1*-INT2 KO).
